# Competition between stacking and divalent cation mediated electrostatic interactions determines the conformations of short DNA sequences

**DOI:** 10.1101/2023.10.05.561104

**Authors:** Balaka Mondal, Debayan Chakraborty, Naoto Hori, Hung T. Nguyen, D. Thirumalai

## Abstract

Interactions between divalent cations (Mg^2+^ and Ca^2+^) and highly charged single stranded DNA (ssDNA) and double stranded DNA (dsDNA), as well as stacking interactions, are important in a variety of problems, including nucleosome stability and phase separation in nucleic acids. Quantitative techniques accounting for ion-DNA interactions are needed to obtain insights into these and related problems. Towards this end, we created a computational model that explicitly takes into account monovalent and divalent ions, within the framework of the sequence-dependent coarse-grained Three Interaction Site (TIS) model for DNA. Molecular simulations of the rigid 24 base-pair (bp) dsDNA and flexible ssDNA sequences, dT_30_ and dA_30_, in a buffer containing Na^+^ and Cl^−^, with varying amounts of the divalent cations, are used to show that the calculated excess number of ions around the dsDNA and ssDNA *agree quantitatively with ion-counting experiments*. Using an ensemble of all-atom structures generated from coarse-grained simulations, we calculated the Small Angle X-ray Scattering (SAXS) profiles, which are also in excellent agreement with experiments. Strikingly, recapitulation of all the experimental findings was achieved without adjusting any of the parameters in the energy function to fit the data. At a molecular level, we find that Mg^2+^ and Ca^2+^ sense the differences between the major and minor grooves in dsDNA even though they are masked in ion-counting and SAXS experiments. The smaller Mg^2+^ binds predominantly to the minor grooves and phosphate groups whereas Ca^2+^ binds specifically only to the minor groove. The dA_30_ conformations are dominated by stacking interactions, resulting in structures with considerable helical order. In contrast, the near cancellation of the favorable stacking and unfavorable electrostatic interactions leads to dT_30_ populating an ensemble of heterogeneous conformations.

## INTRODUCTION

Single and double stranded DNA are polyanions because each phosphate group carries a negative charge. As a result, their conformations are often viewed solely through the lens of electrostatic interactions, which are sensitive to the size, shape and valence of the counterions. Indeed, it is well established that interactions between backbone the phosphate groups and cations, especially multivalent ions, determine the flexibility of DNAs [1]. In particular, cation-mediated changes in the DNA conformations control their propensities to interact with other biomolecules that are involved in several biophysical processes, such as transcription and organization of chromosomes on long length scales [2–4]. For instance, multivalent cations bend even a stiff double stranded DNA (dsDNA) [1, 5]. Ion counting experiments, Small Angle X-ray Scattering (SAXS) measurements, as well as Fluorescence Resonance Energy Transfer (FRET) experiments have provided insights into the interactions of divalent cations with nucleic acids (NAs) [6–10]. Besides electrostatic interactions, the stability of dsDNA should also depend on both Watson-Crick base-pairing (*U*_*HB*_) and base stacking interactions (*U*_*ST*_). The relative contributions of *U*_*HB*_ and *U*_*ST*_, measured in dsDNA by introducing a single nick as a function of temperature and monovalent salt concentration [11] found that the stability of dsDNA was predominantly determined by stacking interactions.

In the 24-base pair (bp) dsDNA (Figure 1A), studied in ion-counting experiments [6], which is used as a case study in this work, Watson-Crick base pairs are fully formed. As a result, the dsDNA is rigid, with the persistence length (≈ 50 nm) being much greater than the contour length (≈ 7.6 nm). Combination of molecular dynamics simulations and integral equation theories successfully reproduced the experimental data from ion counting experiments for a 24-nucleotide dsDNA sequence only in monovalent (NaCl) salt solution [12]. Are there differences between Ca^2+^and Mg^2+^ in the interactions with dsDNA?

**Figure 1:**
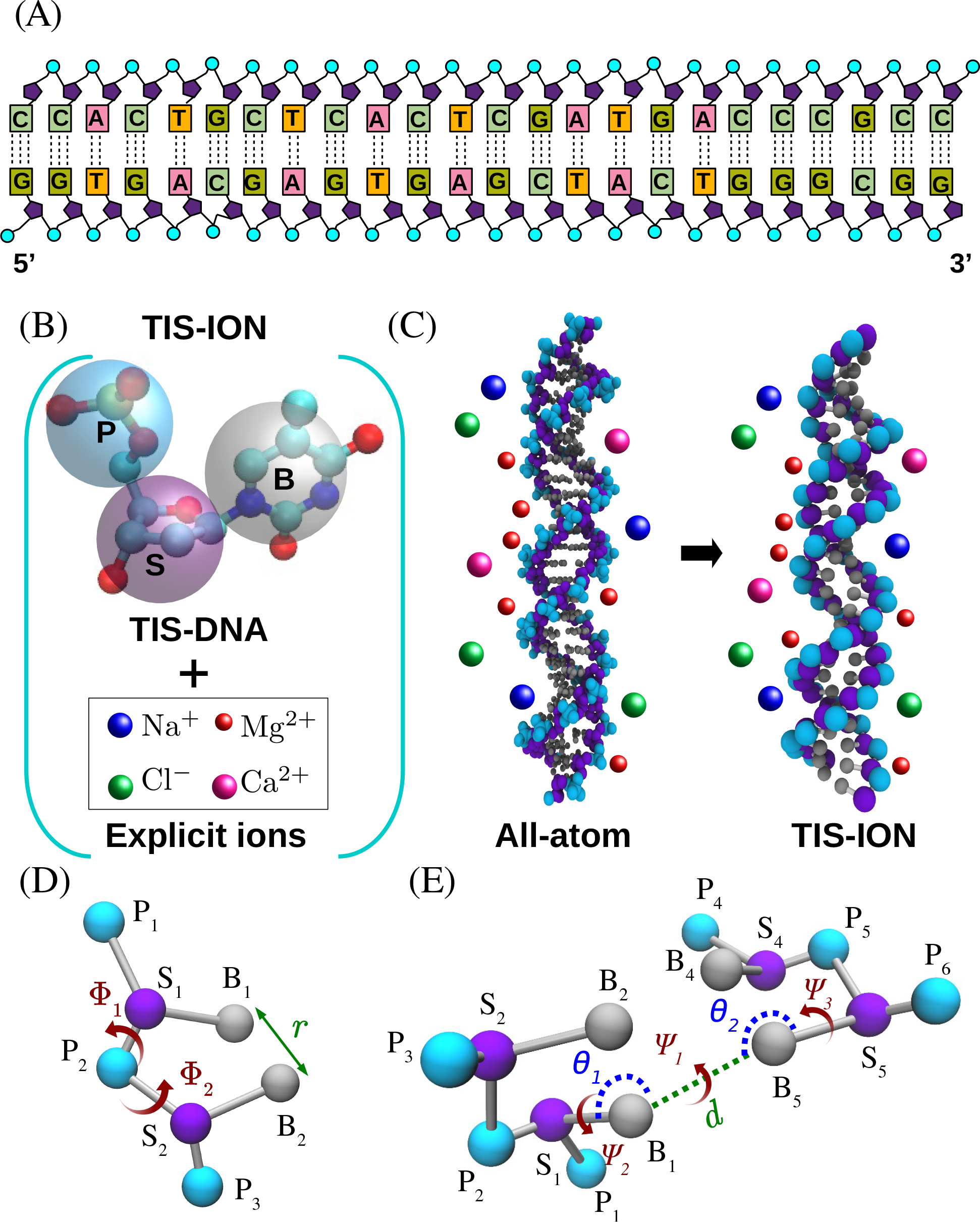
Coarse-grained DNA with explicit ions. (A) Sequence of the 24-bp dsDNA. (B) Coarse-grained representation of a single nucleotide. In TIS-DNA model, each nucleotide is described using three interaction sites, placed at the center of phosphate (P), sugar (S) and base (B) groups, shown in cyan, purple, and grey, respectively. Ions are modeled explicitly using spherical beads of appropriate radii. Na^+^, Mg^2+^, Ca^2+^ and Cl^−^ are shown in blue, red, magenta and green, respectively. (C) All-atom and coarse-grained representations of the dsDNA in Na^+^, Mg^2+^, Ca^2+^ and Cl^−^ ions. Similar color schemes are adapted in all-atom and coarse-grained representations. The three-dimensional (3D) structures of dsDNA were generated using the Nucleic Acid Builder (NAB) [64] software by adopting the parameters for B-DNA. The TIS-DNA [22] coarse-grained structures were generated from the all-atom structures using the visual molecular dynamics (VMD) [65] software. (D) Illustration of the structural features associated with base-stacking (Eq. 3). The distance between adjacent bases B_1_ and B_2_ is *r, ϕ*_1_(P_1_, S_1_, P_2_, S_2_) and *ϕ*_2_(P_3_, S_2_, P_2_, S_1_) are the dihedral angles around sugar-phosphate bonds. (E) Geometry associated with hydrogen bond in Eq. 4. Hydrogen bonding distance *d* is measured between bases B_1_ and B_5_; *θ*_1_(S_1_, B_1_, B_5_) and *θ*_2_(S_5_, B_5_, B_1_) are the angles; *ψ*_1_(S_1_, B_1_, B_5_, S_5_), *ψ*_2_(P_2_, S_1_, B_1_, B_5_) and *ψ*_3_(P_6_, S_5_, B_5_, B_1_) are the dihedral angles.

Ion counting and SAXS experiments of short homopolymeric ssDNAs (poly A and poly T) revealed that their ion-atmospheres are similar [10, 13]. A plausible implication is that the differences in their conformational ensembles must arise from stacking interactions. The brief survey illustrates that experiments cannot resolve the relative importance of *U*_*ST*_, *U*_*HB*_ and ion mediated interactions in modulating the structures of DNA. Single-molecule pulling experiments and theoretical analysis [14, 15] have shown that the bare persistence length of ssDNAs is ≈ 1 nm, which is much smaller than the contour length of even the relatively short sequences (dA_30_ and dT_30_) investigated here. Consequently, cations would affect their conformations to a greater extent than dsDNA, raising the question, how do the conformations of these two ssDNA sequences change as the concentrations of Mg^2+^ and Ca^2+^ are varied? Because purine stacks are preferred over pyrimidine stacks, stacking interactions involving poly A is more favorable than stacking of poly T. Although it is clear that competition of stacking and electrostatic interactions should determine the ensembles of states, a quantitative assessment of their relative contributions has not been made. The brief survey illustrates that experiments alone cannot resolve the relative importance of *U*_*ST*_, *U*_*HB*_ and ion mediated interactions in modulating the structures of DNA. Thus, there is a need to develop well calibrated computational models that not only can take sequence effects into account but also must include key interaction in DNA as well as the role of cations.

Theoretical models, such as, Poisson-Boltzmann (PB) equation [16–19], simulations using Debye-Hückel theory [20–23], or Manning condensation [24] have mimicked the influence of ions on NAs in monovalent salts. However, these models are not as successful in describing DNA interactions with divalent or multivalent cations, where ion-ion correlations are important. Progress has been made in using all-atom (AA) simulations [12, 25–27] to reproduce a few experimental measurements. Excess Na^+^ around the 24-nucleotide dsDNA, calculated using AA simulations in a fixed background of (∼ 5mM) Mg^2+^ [28], is in very good agreement with ion counting experiments. These simulations were performed by keeping the dsDNA rigid, which could be justified because the short sequence is rigid. These studies did not compare the simulation results with experiments (ion counting results and SAXS profiles) over a wide range of divalent ion concentrations. One of the previous coarse-grained (CG) studies, built in the spirit of the TIS model, [29] used surrogate small molecules (dimethylphosphate and magnesium hexahydrate), as done in AA simulations [28] to calculate energetic parameters involving monvalent, divalent, and multivalent cations. The calculated persistent length for dA_68_ as a function of Na^+^ is in reasonable agreement with experiments. None of the models for dsDNA and ssDNA have simultaneously reproduced a variety experimental measurements for both monovalent and divalent cations over a broad concentration range, a deficiency that has to be overcome.

In order to overcome the difficulties associated with the previous models, we created a new coarse-grained model that is based on the TIS model for NAs [30, 31]. Each nucleotide is represented as three spherical beads that are located at the centers of mass of phosphate (P, in cyan), sugar (S, in purple) and base (B, in grey) groups (Figure 1B). In some of our previous studies on NAs, electrostatic interactions between the phosphate (P) groups were modeled using the Debye-Hückel potential with a reduced charge on P in order to account for counter ion condensation [21, 22]. Although the TIS model, with the implicit treatment of ions, is successful and is the basis on which other three site models for NAs were proposed [32–34], the lack of explicit inclusion of monovalent and divalent cations makes it impossible to investigate myriad of problems involving DNA biophysics. Here, we combine the TIS model for DNA [22] and explicit treatment of monovalent and divalent cations to create the TIS-ION model. To develop the new model, we followed the strategy used in the context of RNA folding [35–37] in which the effects of Mg^2+^, and Ca^2+^, monovalent cations were modeled explicitly. The resulting TIS-ION model, which also accounts for sequence effects in DNA, is a significant advance, as we demonstrate here by quantitative comparison with experiments, *without using a single adjustable parameter* to fit any of the data. The simulations quantitatively reproduce the experimental findings for the dependence of the number of ions around short DNA constructs as a function of [Mg^2+^]. The corresponding results for Ca^2+^ are predictions, which can be readily tested. We also find that the agreement between the calculated and experimental SAXS profiles, for a 25-nucleotide dsDNA, dT_30_ and dA_30_ are excellent. Strikingly, simulations show that in these short DNA sequences (dsDNA and dA_30_), the conformations are determined by stacking interactions, with electrostatic interactions playing a minor role.

## METHODS

### TIS-ION model

The energy function in the TIS-ION model is,

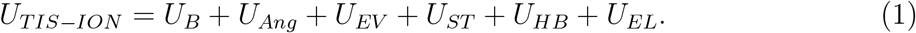

The potentials for bond stretch (*U*_*B*_) and bond angle (*U*_*Ang*_) are given by, *U*_*B*_ = *k*_*b*_(*r* − *r*_0_)^2^ and *U*_*Ang*_ = *k*_*θ*_(*θ* − *θ*_0_)^2^, where *r*_0_ and *θ*_0_ are the equilibrium bond length and bond angle, respectively, which were taken from an ideal B-form DNA helix. The values of *k*_*b*_, *k*_*θ*_, *r*_0_, and *θ*_0_, are listed elsewhere [22]. Excluded volume repulsion between sites *i* and *j* (DNA sites or ions) separated by a distance, *r* (in nm), is modeled as,

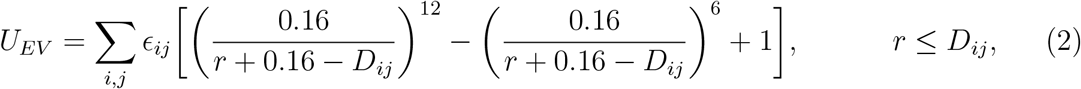

where *D*_*ij*_ = *R*_*i*_ + *R*_*j*_ and 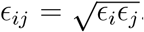. The values of radius *R* and *ϵ* for the ions and DNA are listed in Table S1 of the Supporting Information (SI). When both the interacting sites belong to DNA, *D*_*ij*_ is taken to be 0.32 nm [21].

### Stacking Potential

The potential for stacking interactions, *U*_*ST*_, between two consecutive nucleotides, *i* and *i* + 1, is taken to be [21, 22],

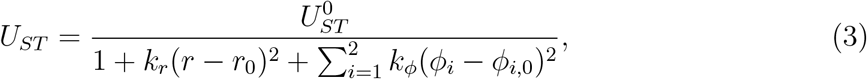

where *r* is the distance between the bases B_i_ and B_i+1_, and *ϕ*_1_ (P_i_, S_i_, P_i+1_, S_i+1_) and *ϕ*_2_ (P_i+2_, S_i+1_, P_i+1_, S_i_) are the dihedral angles around the sugar-phosphate bonds (see Figure 1D). The equilibrium stacking distance *r*_0_, and the backbone dihedral angles *ϕ*_1,0_ and *ϕ*_2,0_ are calculated from the coarse-grained representation of an ideal B-form DNA helix.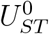 gives an estimate of stacking energy between two consecutive nucleotides in an ideal B-form DNA helix, and is ≈ -4.19 kcal/mol for a T-dimer and -5.77 kcal/mol for an A-dimer, at T = 277 K [22]. It is worth noting that the 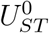 values are not fit parameters, but calibrated in our previous work [22] to accurately reproduce the thermodynamics of DNA duplexes, following the nearest neighbor model [38, 39].

### Hydrogen bond interactions, *U*_*HB*_

Hydrogen bonds are considered between canonical base pairs (Watson-Crick), belonging to the complementary strands. The *U*_*HB*_ potential is,

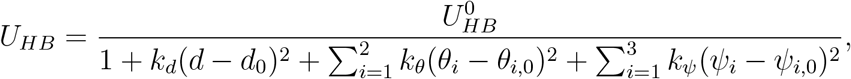

where *d, θ*_1_, *θ*_2_, *ψ*_1_, *ψ*_2_ and *ψ*_3_ are described in Figure 1E. The parameters in *U*_*HB*_, along with the procedures used to evaluate them, are reported in our previous study [22]. Hydrogen bond interactions are absent in the homopolymeric ssDNA sequences.

### Electrostatic interactions, *U*_*EL*_

The interaction between two charged groups with charges *Q*_*i*_ and *Q*_*j*_, separated by a distance, *r*_*ij*_, is given by the Coulomb potential,

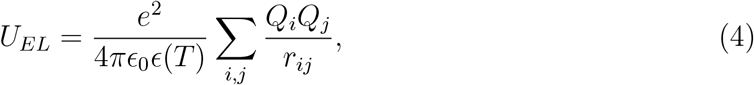

where *ϵ*_0_ is the vacuum permittivity, and the temperature-dependent dielectric constant, *ϵ*(*T*) = 87.74 − 0.4008T + 9.398 × 10^−4^*T* ^2^ − 1.410 × 10^−6^*T* ^3^, is taken from [40], where the temperature, *T*, is reported in celcius. The charge on the phosphate group is *Q*_*P*_ = −1e (unit of electron charge).

### Simulations

We used OpenMM [41], a high-performance toolkit for molecular simulations, to simulate ssDNA and dsDNA in mixed salt solutions of monovalent and divalent cations, on a Graphics Processing Unit (GPU). We used the Particle-Mesh-Ewald (PME) algorithm [42], as implemented in OpenMM to account for the long-range Coulomb interactions. In order to enhance conformational sampling, the simulations were performed by integrating the Langevin equation in the low friction limit [43]. The equation of motion for a bead or ion, with coordinate 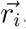, is,

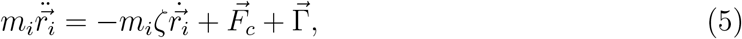

where *m*_*i*_ is the mass of the bead, *ζ* is the friction coefficient, 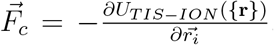, and 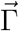 is the random force with a white noise spectrum. The autocorrelation function of the random force in the discretized form is 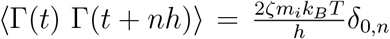, where *n* = 0, 1, … and *δ*_0,n_ is the Kronecker delta function. The equations of motion were integrated using Langevin leap-frog method [44], as implemented in OpenMM, using *ζ*= 0.01 picosecond^−1^. The integration time step is 2 femtoseconds. We performed simulations at *T* = 277 K to compare the results with the ion-counting experiments [6, 10]. Additional simulations were performed at *T* = 298 K to compare with the SAXS experiments [7, 45].

The systems consist of a single DNA in a cubic box of length L whose value depends on the ion concentration. To minimize finite size effects, we used periodic boundary conditions. Neutrality of the sample is maintained by adding an appropriate number of Cl^−^ ions.

### Counting ions in the vicinity of Nas

The experimentally measurable Γ_*i*_, which is an estimate of the excess number of cations *i*, in the vicinity of NAs, is calculated using the Kirkwood-Buff integral [46, 47],

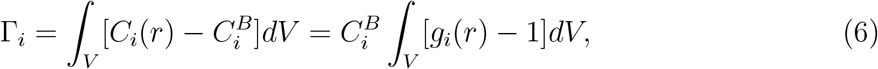

where V is the volume over which the integral is evaluated. The concentration of ions *i* in the vicinity of the polyanion, 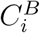, differs from the bulk concentration 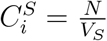, where *N* is the total number of ions *i*, and *V*_*S*_ (=*L*^3^) is the volume of the simulation box. We calculated 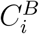 for the *i*^th^ ionic species after the concentration profile reaches a plateau at large separation from the DNA (see Supplementary Information (SI), Figure S1). In practice, the integral in Eq. 6 is carried out over a finite volume because the density fluctuations are predominantly localized near the NA surface.

For dsDNA, the volume integral is,

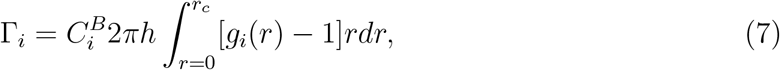

where *h* is the approximate height of the cylinder encompassing the dsDNA along the z axis (helical axis or the principal axis), *r* is the radial distance from the principal axis of the dsDNA and 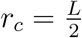, *L* is the simulation box length (see Figure S2). We varied *h* to obtain convergence in the Γ_*i*_ values and chose 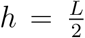 (see Figure S3). The size of the smallest simulation box is *L* ≈ 23 nm which is significantly larger than the length of dsDNA ≈ 7.6 nm. For ssDNA, the volume integral is,

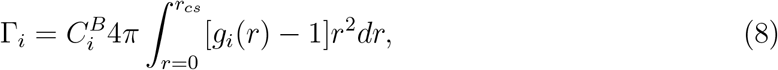

where 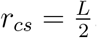, and *r* is the distance of the ion *i* from the center of mass of ssDNA (Figure S2B in the SI).

### Calculation of Γ_*i*_ from simulations

We performed Langevin dynamics simulations of the 24 bp dsDNA and ssDNA dA_30_ and dT_30_, in a solution containing both monovalent and divalent ions (*T* = 277 K). First, we performed simulations for each DNA construct at different concentration of Mg^2+^ or Ca^2+^ by varying the box length from ≈ 23 nm to 83 nm. We used a constant Na^+^ concentration (= 20 mM), and varied [Mg^2+^] or [Ca^2+^] from 0.1 mM to 20 mM (in case of ssDNA) or 50 mM (in case of dsDNA) to closely mimic the experiments [6, 10]. We calculated the Γ_*i*_ values for each ionic species by averaging over all the sampled conformations after equilibrium has been reached using either Eq. 7 (dsDNA) or Eq. 8 (ssDNA).

### Stacking in ssDNA

We describe stacking in ssDNA by introducing an order parameter, *p*_*S*_ – a measure that is related to the helix content [48] in a biomolecule. We assume that each nucleotide is either in a stacked (S) state or in a random coil state (C). To measure the stacking propensity of the *i*^*th*^ nucleotide, we calculated the sugar-phosphate dihedral angles, *ψ*(*P*_*i*_, *S*_*i*_, *P*_*i*+1_, *S*_*i*+1_) and *ϕ*(*S*_*i*_, *P*_*i*+1_, *S*_*i*+1_, *P*_*i*+2_), and compared them with the typical *ψ*_*B*_ and *ϕ*_*B*_ values in an ideal B-form helix. If the deviations *δψ*(= |*ψ* − *ψ*_*B*_|) and *δϕ*(= |*ϕ* − *ϕ*_*B*_|) are within 25% of the ideal helix parameters, we consider the nucleotide to be in the S state. For a given ssDNA conformation, *p*_*S*_ is defined as,

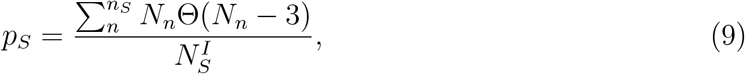

where *n*_*S*_ is the number of consecutive stacks along the ssDNA contour, *N*_*n*_ is the length of the *n*^*th*^ stack and 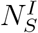 is the length of the persistent stacks in completely stacked ideal B helix and Θ(*N*_*n*_ − 3) is unity only if *N*_*n*_ ≥ 3. A minimum of three consecutive nucleotides (*N*_*n*_ ≥ 3) should be in the state *S* to calculate *p*_*S*_. For instance, in the oligomer 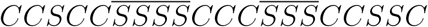, there are two persistent stacks (*n*_*S*_ = 2) of length 4 and 3, (*N*_1_ = 4, *N*_2_ = 3) respectively, yielding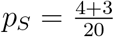, which means that there are 7 nucleotides that are persistently stacked. In the example, the lengths of the longest stretch of stacked and unstacked bases are, 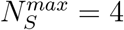 and 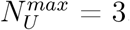, respectively.

## RESULTS

### Comparison with ion counting experiments

We compare in Figures 2A and B the calculated and experimental [6] 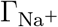 (blue) and 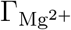 (red) or 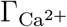 (magenta) for dsDNA as a function of [Mg^2+^] (or [Ca^2+^]). The agreement is excellent for both monovalent and divalent cations over the entire concentration range. Because multivalent cations are more effective in screening the phosphate charges, monovalent cations (Na^+^) are replaced by divalent cations (X^2+^) as the concentration of Mg^2+^ or Ca^2+^ is increased. This is a consequence of the entropically favored counterion release mechanism [49–53].

**Figure 2:**
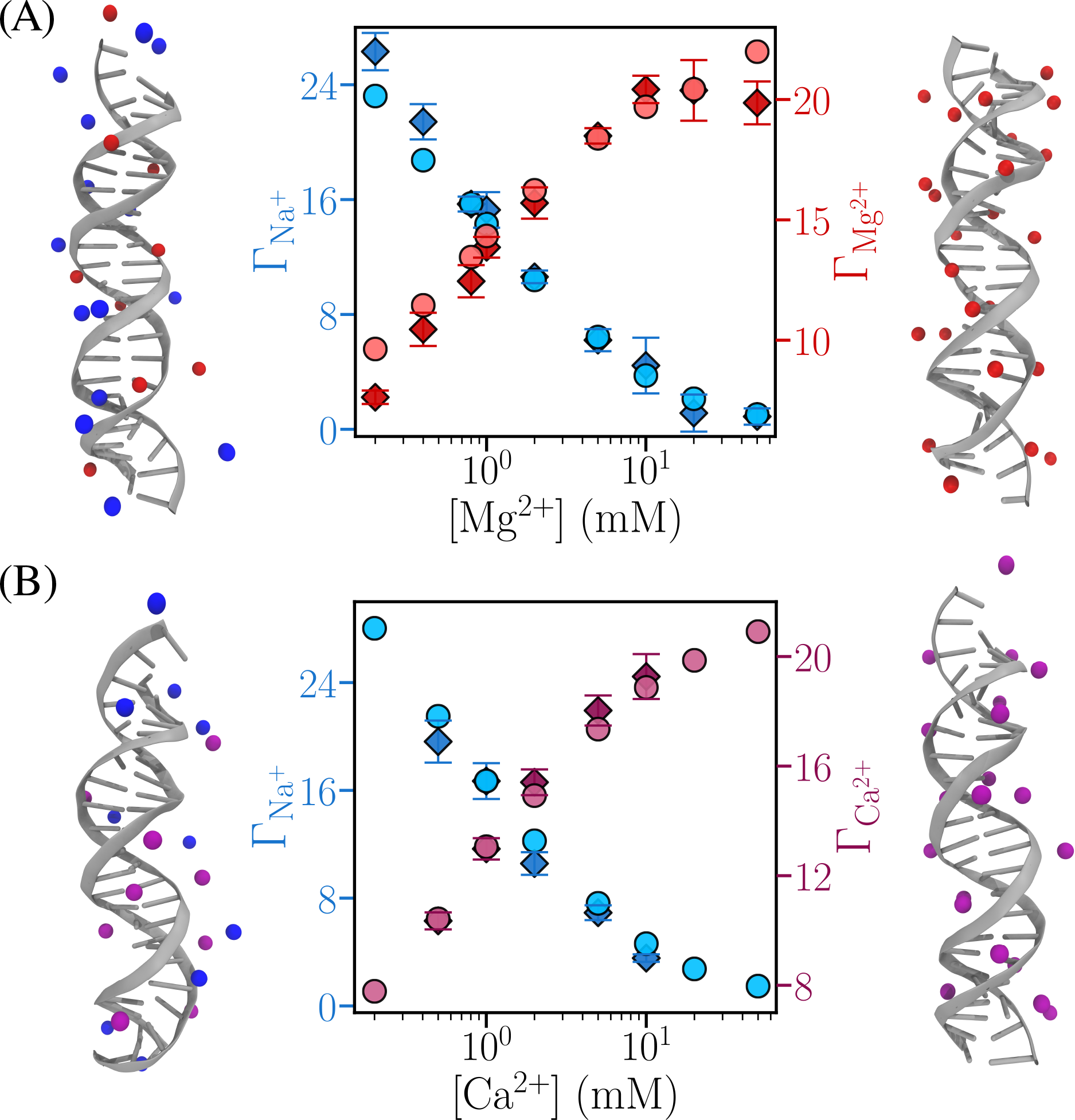
Ions around dsDNA. (A) Excess number of ions, 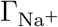 (blue) and 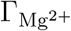 (red), in the vicinity of dsDNA as a function of [Mg^2+^]. Experimental results [6] in diamonds in dark shade, and the simulation results are in circles in light shade. Reconstructed all-atom structures, highlighting Na^+^ and Mg^2+^ ions within 10 Å of the phosphate groups, are shown for [Mg^2+^] = 0.2 mM and 20 mM. (B) Same as (A), except the results are shown for Ca^2+^ ions (in magenta). In all the panels, [Na^+^] = 20 mM.

The simulation results for Γ_*i*_, as a function of [Mg^2+^], for dT_30_ and dA_30_. is also in very good agreement with experiments [10] for both dT_30_ and dA_30_ (Figure 3). Interestingly, there is negligible difference in Γ_*i*_ for dT_30_ and dA_30_, which implies that the preferential ion coefficient cannot be used to determine sequence effects, at least in short homopolymeric ssDNA sequences. As demonstrated below, notable distinctions exist in the conformational characteristics of dT_30_ and dA_30_.

**Figure 3:**
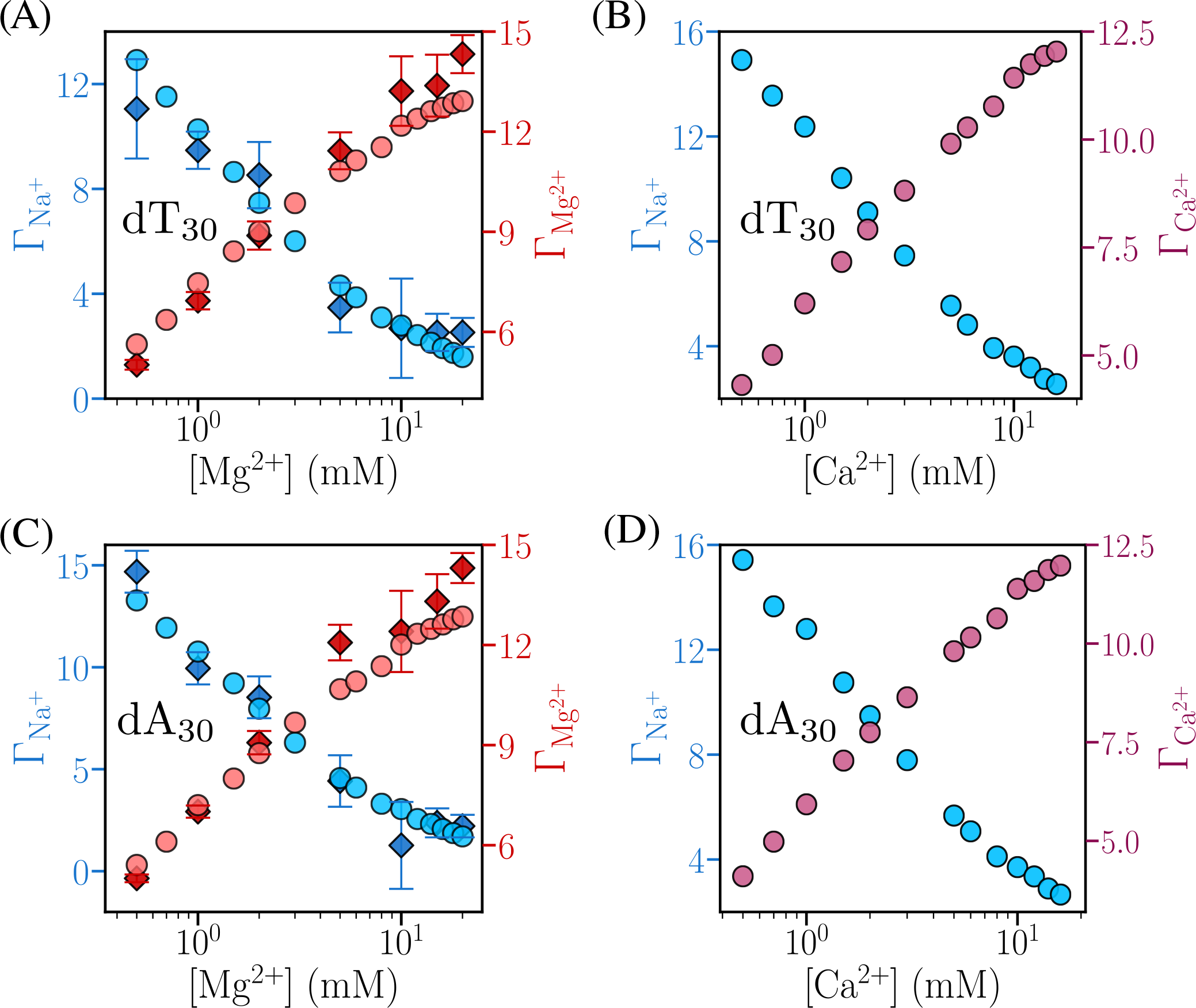
Ions around ssDNA. (A) Excess number of ions, 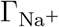 (in blue) and 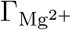 (in red), in the vicinity of ssDNA sequence dT_30_ as a function of [Mg^2+^]. Experimental results [10] are in the dark shade diamonds. Simulation results are in circles in light shade. (B) Same as (A), except the results are for Ca^2+^ ions (in magenta). (C) Same as (A), except results are for the dA_30_ sequence. (D) Same as (C), except results are for Ca^2+^ ions. [Na^+^] = 20 mM in the panels.

Comparison of the results for Mg^2+^ localization around dsDNA, dT_30_ and dA_30_ allows us to infer the following observations: (1) The Mg^2+^ concentration, at which Na^+^ is released is ≈ 1 mM for dsDNA, whereas it is ≈ 2.5 mM for dT_30_ and dA_30_. This is likely due to the large difference in flexibility between dsDNA and ssDNA. (2) Interestingly, the number of cations around dT_30_ and dA_30_, as a function of [Mg^2+^] is almost the same although there is a significant difference in their conformational fluctuations (see below). (3) Somewhat surprisingly, our simulations as well as experiments [6], predict that 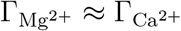, at all values of the divalent ion concentration even though their charge densities are different.

### Agreement between calculated and experimental SAXS profiles is excellent

We calculated the SAXS profiles, *I*(*q*), for a 25 bp dsDNA (see Figure 4A for the sequence), dT_30_ and dA_30_ at *T* = 298 K, as a function of the wave vector *q*, using simulations in order to compare with experiments [7, 10, 45]. The experimental SAXS profiles [10] were accessed either from the publicly available SAXS database SASDBD (dT_30_: SASDBD6, dA_30_: SASDBE6, in 20 mM NaCl solution) or extracted using the WebPlotDigitizer [54] from the published data (dT_30_ in salt solution containing 20 mM Mg^2+^ and 20 mM NaCl [7], dsDNA in 3 mM and 16 mM Mg^2+^, and 0.4 mM NaCl [45]). We calculated *I*(*q*) using at least 5,000 randomly chosen conformations using the TIS2AA software [55] and CRYSOL [56]. Before calculating *I*(*q*), we converted the structures generated in the coarse-grained simulations to all-atom conformations using an in-house code. The resulting ensemble of all-atom structures proves to be convenient in the calculation of SAXS profiles.

**Figure 4:**
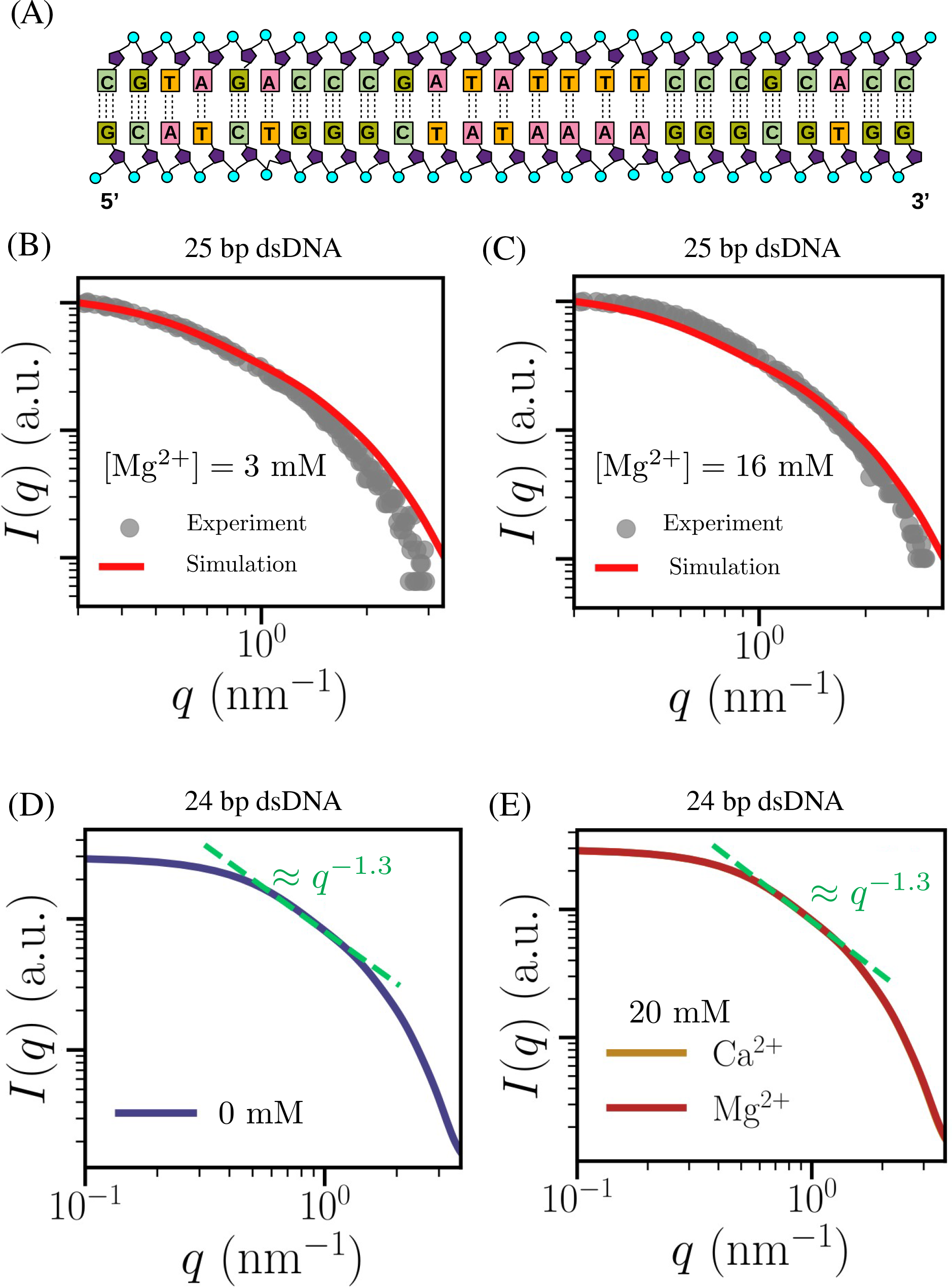
SAXS profiles for dsDNA: (A) Sequence of the 25-bp dsDNA. (B) Comparison of experimental [45] and simulated SAXS profiles in [Mg^2+^] = 3 mM and [Na^+^] = 0.4 mM at *T* = 298 K. (C) Same as (B), except the results are for [Mg^2+^] = 16 mM. (D) SAXS profile of the 24 bp dsDNA (Figure 1(A)) in [Na^+^] = 20 mM at *T* = 277 K. (E) Comparison of SAXS profiles of the same dsDNA sequence, in [Mg^2+^] = 20 mM and [Ca^2+^] = 20 mM, in [Na^+^] = 20 mM.

### dsDNA

We first calculated *I*(*q*) as a function of *q* at 3 mM and 16 mM [Mg^2+^], for the 25 bp dsDNA for which experimental [45] data is available. The results in Figures 4B and 4C show that the simulations reproduce the experimental results almost quantitatively. There are only minor deviations at *q* > 2 nm^−1^. Few of the reconstructed all-atom structures used to calculate the SAXS profiles for the 25 bp dsDNA are shown in Figure S4 in the SI.

Measurements of *I*(*q*) are not available for the dsDNA sequence studied in the ion counting experiments. Nevertheless, the accuracy of our simulations allows us to predict the outcomes and compare the results for Mg^2+^ and Ca^2+^. From Figures 4D and 4E, which show *I*(*q*) as a function of *q* at 0 mM and 20 mM divalent cation concentrations, it is clear that the SAXS profiles are almost identical for both the ions. The length of the dsDNA (24 bp) is ≈ 6 times smaller than the bare persistence length of a typical dsDNA (150 bp). The dominance of the bare persistence length in the short dsDNA, due to WC base pairing, implies that the 24-nucleotide sequence should be impervious to cations, thus explaining the *I*(*q*) results in Figure 4. The dependence of *I*(*q*) ∼ *q*^−1.3^ (see below) at the intermediate *q* values (Figure 4D, 4E), suggests that the dsDNA is not a rod, but behaves like a stiff polymer [57].

### SAXS profiles for ssDNAs show scale-dependent structural changes

Figure 5, with comparisons to experiments in the insets, displays the calculated SAXS profiles for dT_30_ and dA_30_. The simulated and the experimental SAXS profiles are in excellent agreement. The variations of *I*(*q*) with *q* reveals details about the changes in the structure of the ssDNA sequences at different length scales. Scaling relations of *I*(*q*) with *q* are known for various polymer shapes. For example, *I*(*q*) ∝ *q*^−*x*^, where *x* = 2 (Gaussian chains), *x* = 1 (rods), *x* = 5/3 (polymer in good solvent), and *x* = 4 (for globules) [57]. At [X^2+^] = 0 mM, in the range *q* ∼ 0.8 − 1.5 nm^−1^, *x* ≈ 1.4 for dT_30_. In contrast, for dA_30_, *x* ≈ 1.6, which is close to the value expected for polymers in good solvents. The value of *x* increases for dT_30_ (*x* ≈ 1.8), which implies that there is change in conformation from a stiff to more flexible polymer behavior as [X^2+^] increases. In contrast, *x* ≈ 1.6 is roughly the same for dA_30_ at the elevated [X^2+^]. Examples of the reconstructed all-atom structures, used to calculate SAXS profiles for dT_30_ and dA_30_, are shown in Figures 5E, S5 and S6, respectively.

**Figure 5:**
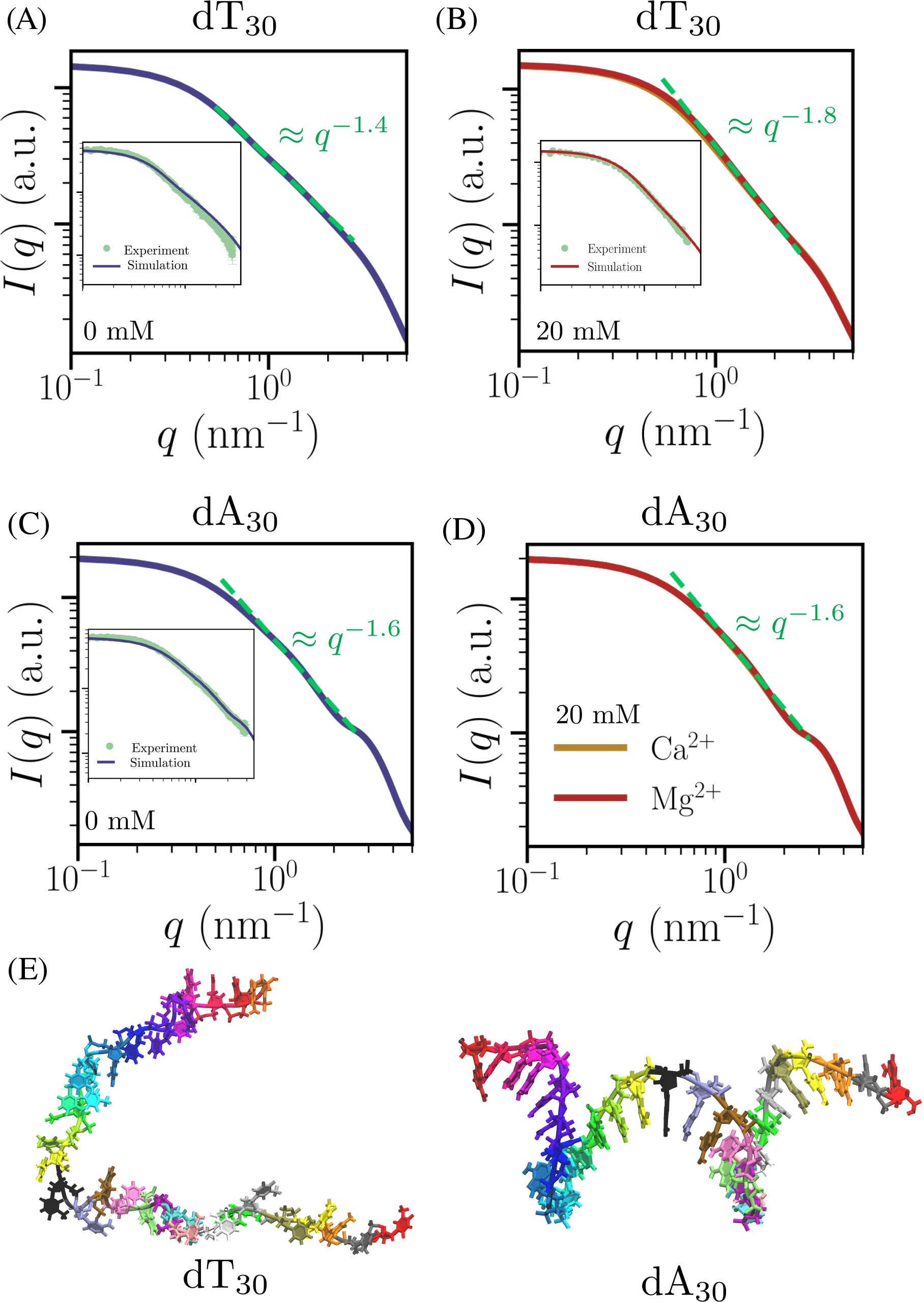
Comparison of simulated and experimental SAXS profiles for ss-DNA: (A-D) SAXS profiles are computed for dT_30_ and dA_30_, in [Mg^2+^] or [Ca^2+^], at 0 mM and 20 mM, in [Na^+^] = 20 mM at *T* = 277 K. Inset compares the simulated and experimental SAXS profiles with error bars, when available, at *T* = 298 K. (E) Representative all-atom structures, reconstructed from coarse-grained simulations, are shown at [Mg^2+^] = 20 mM. To aid visualization, nucleotides are assigned different colors based on their sequence number.

### Stacking versus electrostatic interactions

It can be argued that the differences in *x* values in dT_30_ and dA_30_ as [X^2+^] increases should be a consequence of the interplay between favorable stacking and cation-mediated electrostatic repulsion between the phosphate groups. In dA_30_ the stacking interactions dominate, and essentially determine the conformations of the ssDNA. Indeed, the favorable stacking interaction (⟨*U*_*ST*_ ⟩), is more stabilizing than the effective repulsion between the phosphate groups 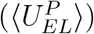 in dA_30_ (see Figure S7 in SI). Thus, ⟨*U*_*ST*_ ⟩, with 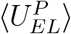 playing a minor role, determines the structural ensemble of dA_30_, which explains the constancy of *x* with increasing [Mg^2+^]. In contrast, the values of ⟨*U*_*ST*_ ⟩ and 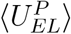 are comparable in dT_30_. As a result, thermal fluctuations play an important role in shaping the conformations of dT_30_. At low divalent concentrations, there is a preference to populate stiff conformations, which become more flexible at the highest divalent concentrations. The enhanced preference for stacking in Adenine compared to Thymine rationalizes the simulation results. A natural prediction is that balance between ⟨*U*_*ST*_ ⟩ and 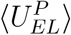 may be altered in dT_30_ at lower temperatures.

### Distribution of *R*_*g*_

The results in Figure 3 show that there are little variations in 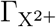 as a function of [X^2+^], despite the known differences on the stacking interactions between A and T [9]. The sequence-dependent variations are reflected in the distributions *P*(*R*_*g*_) of the radius of gyration, *R*_*g*_ (Figures 6A-D). The dispersion in *P*(*R*_*g*_) changes considerably as [Mg^2+^] or [Ca^2+^] is increased. At [X^2+^] = 20 mM, dT_30_ samples a broad range of conformations (Figures 6A and 6B). Although qualitatively similar results are found for dA_30_ (Figures 6C and 6D), the changes in the width of *P*(*R*_*g*_) with increase in [X^2+^] is less pronounced.

**Figure 6:**
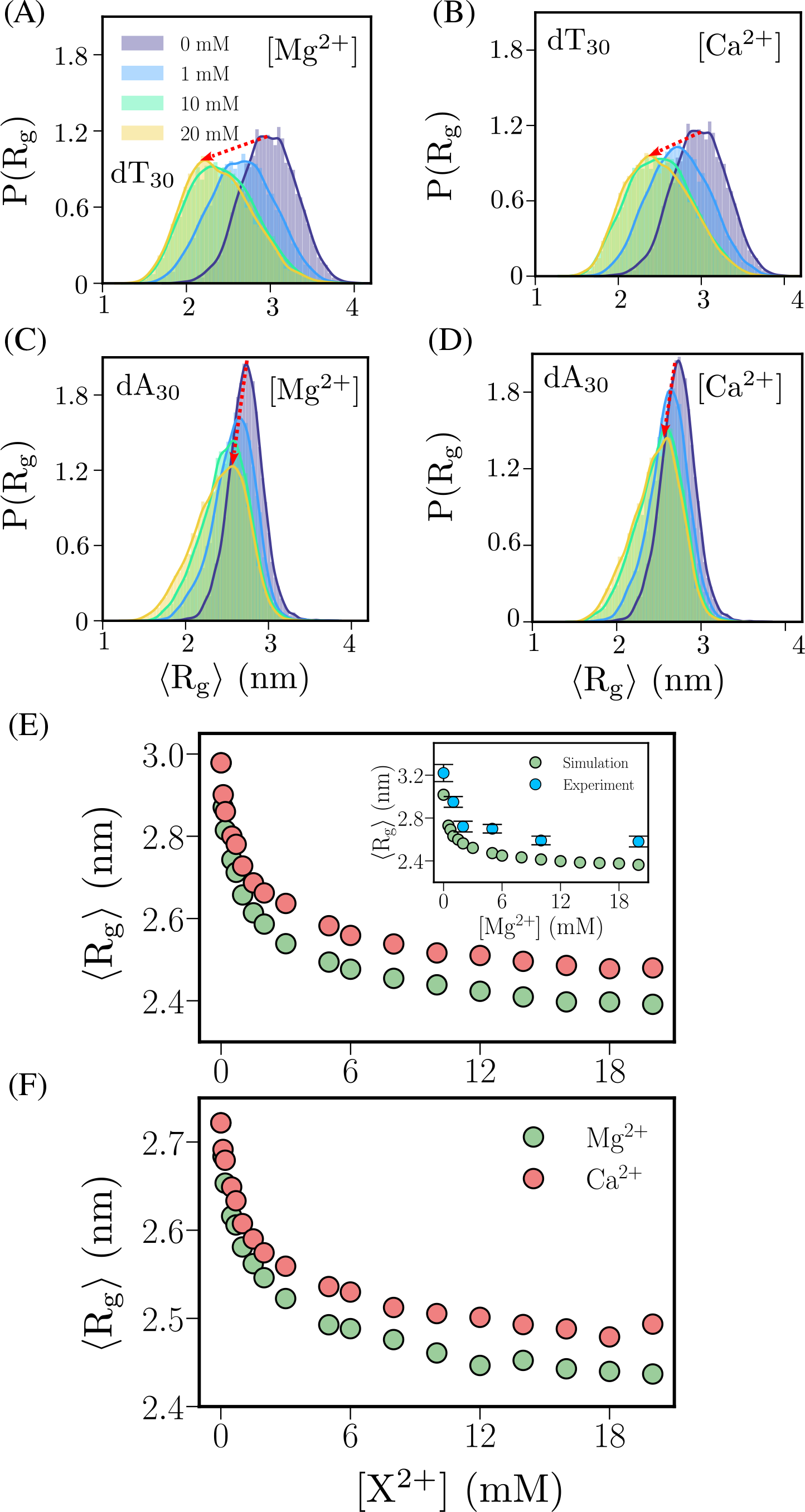
[X^2+^] dependent changes in the radius of gyration: (A-D) Normalized distribution of *R*_*g*_ for dT_30_ and dA_30_, as a function of [Mg^2+^] and [Ca^2+^]. Movement of the peak of the distributions are indicated with arrows. (E) Mean value, ⟨*R*_*g*_⟩ of dT_30_ as a function of [X^2+^], where X^2+^ = Mg^2+^ or Ca^2+^, in [Na^+^] = 20 mM at *T* = 277 K. Inset compares the calculated and experimentally measured [7] ⟨*R*_*g*_⟩ for dT_30_ at 298 K in 20 mM NaCl solution. (F) Same as (E), except the results are for dA_30_ sequence.

Comparison of the calculated and extracted mean ⟨*R*_*g*_⟩ from experimental SAXS profiles [7] for dT_30_ at *T* = 298 K (see inset in Figure 6E), shows very good agreement at all values of [X^2+^]. Although the simulations predict a somewhat larger degree of compaction, the differences between experiments and simulations are small. The ⟨*R*_*g*_⟩ values in Ca^2+^ are higher than in Mg^2+^ for both dT_30_ and dA_30_ (Figures 6E and 6F), which in effect means that Mg^2+^ induces more compaction because of the higher charge density.

### Effective Flory exponent

In order to elucidate the polymer properties of the ssDNAs, we fit the calculated mean distance *R*_*ij*_ between the phosphate groups *i* and *j*, to the equation,

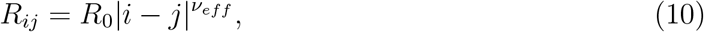

where *R*_0_ is the prefactor, and *v*_*eff*_ is the effective Flory exponent. Because of the limited range of |*i* − *j*| (Figure 7), the extracted values of *v*_*eff*_ and *R*_0_ might not be accurate. Nevertheless, Figures 7A-7D show that ln *R*_*ij*_ ≈ *v*_*eff*_ ln |*i* − *j*| at all [X^2+^] is linear, thus allowing to extract *v*_*eff*_ as a function of [Mg^2+^] (Figure 7E) and [Ca^2+^] (Figure 7F). The plots clearly show the influence of electrostatic interactions.

**Figure7:**
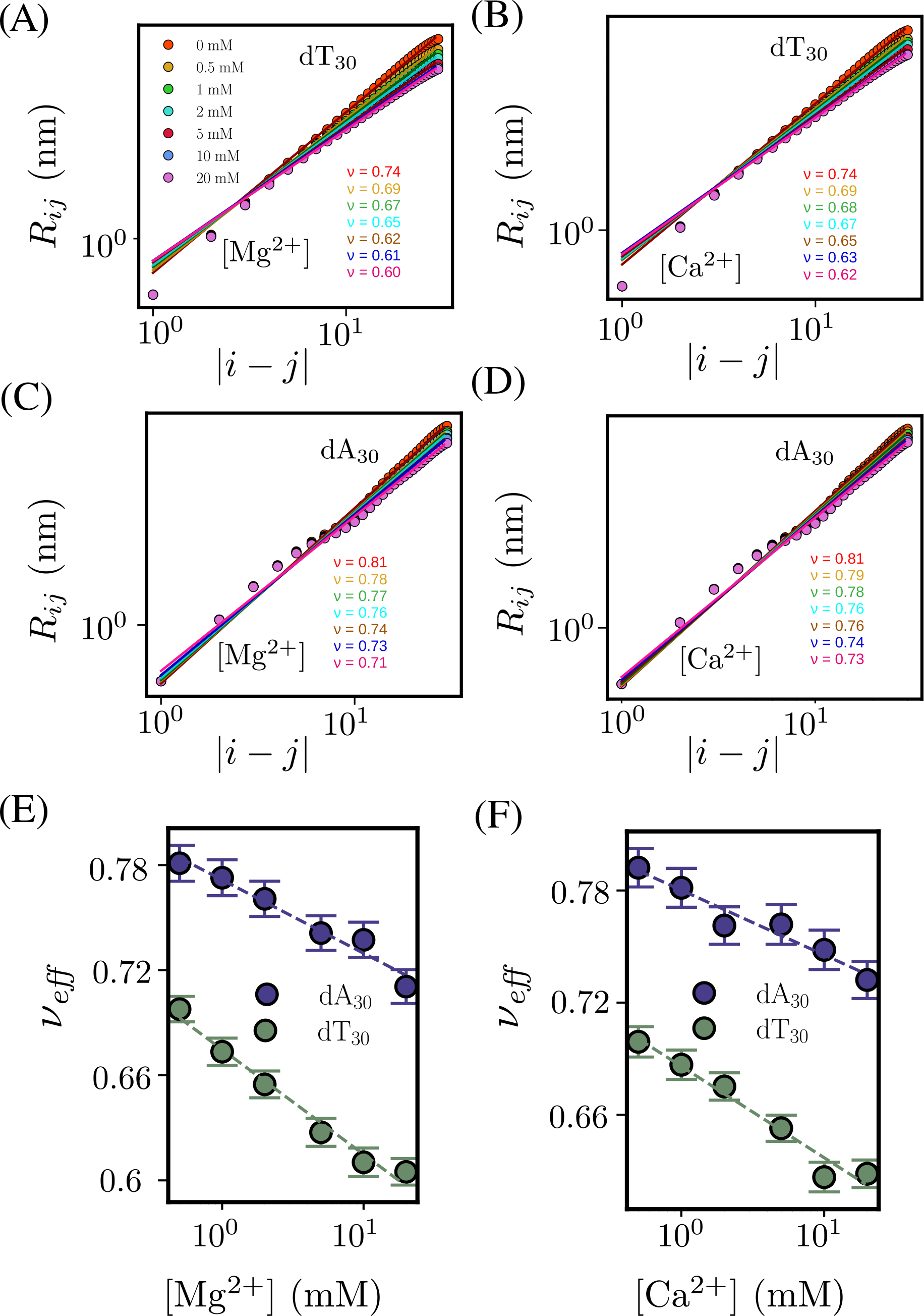
Effective Flory exponent: (A) Distance between the phosphate groups of dT_30_, *R*_*ij*_, as a function of nucleotide separation, |*i* − *j*|, along the contour, at different [Mg^2+^], in [Na^+^] = 20 mM. (B) Same as (A), except the results are for [Ca^2+^]. (C) Same as (A), except the results are for dA_30_ sequence. (D) Same as (C), except the results are for [Ca^2+^]. (E) Effective Flory exponent, *v*_*eff*_ as a function of [Mg^2+^], for dT_30_ and dA_30_. (F) Same as (E), except the results are for [Ca^2+^].

A few observations pertaining to the internal structures follow from Figures 7E and 7F. (1) The values of *v*_*eff*_ decrease linearly as [X^2+^] increases in both the sequences. At all [X^2+^], *v*_*eff*_ for dA_30_ is significantly larger than dT_30_ in both the divalent cations. (2) At higher [X^2+^] (= 20 mM), *v*_*eff*_ ≈ 0.6 for dT_30_, which is the characteristic value for a homopolymer in a good solvent. Interestingly, only at ≈ 1 M monovalent salt concentration, *v*_*eff*_ ≈ 0.6 [22, 58] for poly T. (3) Because *v*_*eff*_ for dA_30_ exceeds 0.6 at all [X^2+^], it follows that poly A sequences are likely to behave as semiflexible chains. The stiffness of dA_30_ can be attributed to the propensity of adenosine bases to form stable stacks. This is consistent with the minimal changes observed in SAXS scaling relations (*x* = 1.6) for dA_30_, as divalent cation concentrations are increased from 0 to 20 mM. (4) The prefactor *R*_0_ for dT_30_ in Mg^2+^ varies from 0.7 nm - 0.8 nm in the concentration range (0 - 20) mM (Table S2 in the SI). The values of *R*_0_ ≈ (0.5 - 0.6) nm is smaller for dA_30_ (Table S2 in SI). The prefactor is a measure of the mean distance between the nucleobases. The smaller value of *R*_0_ in dA_30_ is due to the more favorable stacking interactions in Adenine compared to Thymine.

### Stacking interactions stiffen dA_30_

The procedure used to calculate ⟨*p*_*S*_⟩ is illustrated in Figures 8A and 8B. The distributions of the longest stacked 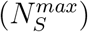 and unstacked 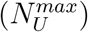 bases in [Mg^2+^] = 20 mM, are shown in Figures 8C and 8D, respectively. For dT_30_, 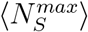 ≈ 4 nucleotides (nt), whereas for dA_30_, the peak appears at 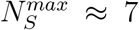 nt, with a tail that extends to 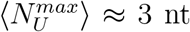. The longest unstacked segments in dA_30_ has a narrower distribution with 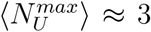 nt compared to dT_30_, for which the distribution extends to ≈ 16 nt. Long helical stacks are interrupted by short unstacked bases in dA_30_, whereas shorter helical stacks are interrupted by long-stretches of unstacked segments in dT_30_. For instance, the probability of finding dT_30_ conformations with 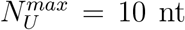, which is the periodicity of a B-DNA helix, is ≈ 5 % whereas it is only 0.02 % (a decrease by a factor of 250) in dA_30_. Figure S8 compares the distribution 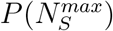 and 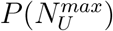 at [Mg^2+^] = 0 mM and 20 mM, in dT_30_ and dA_30_. Figures 8E and 8F show that 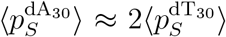 at all [X^2+^], and is independent of the nature of the cation. This finding explains the increased stiffness of dA_30_ relative to dT_30_, which accords well with the observation that Adenine bases participate in stronger base stacking compared to Thymine [7, 9, 10, 59]. The independence of ⟨*p*_*S*_⟩ on [X^2+^] finds support in the magnetic tweezer experiments [9] that report nearly constant free energy difference between stacked poly(dA) and unstacked polypyrimidine over three orders of magnitude in NaCl concentration.

**Figure 8:**
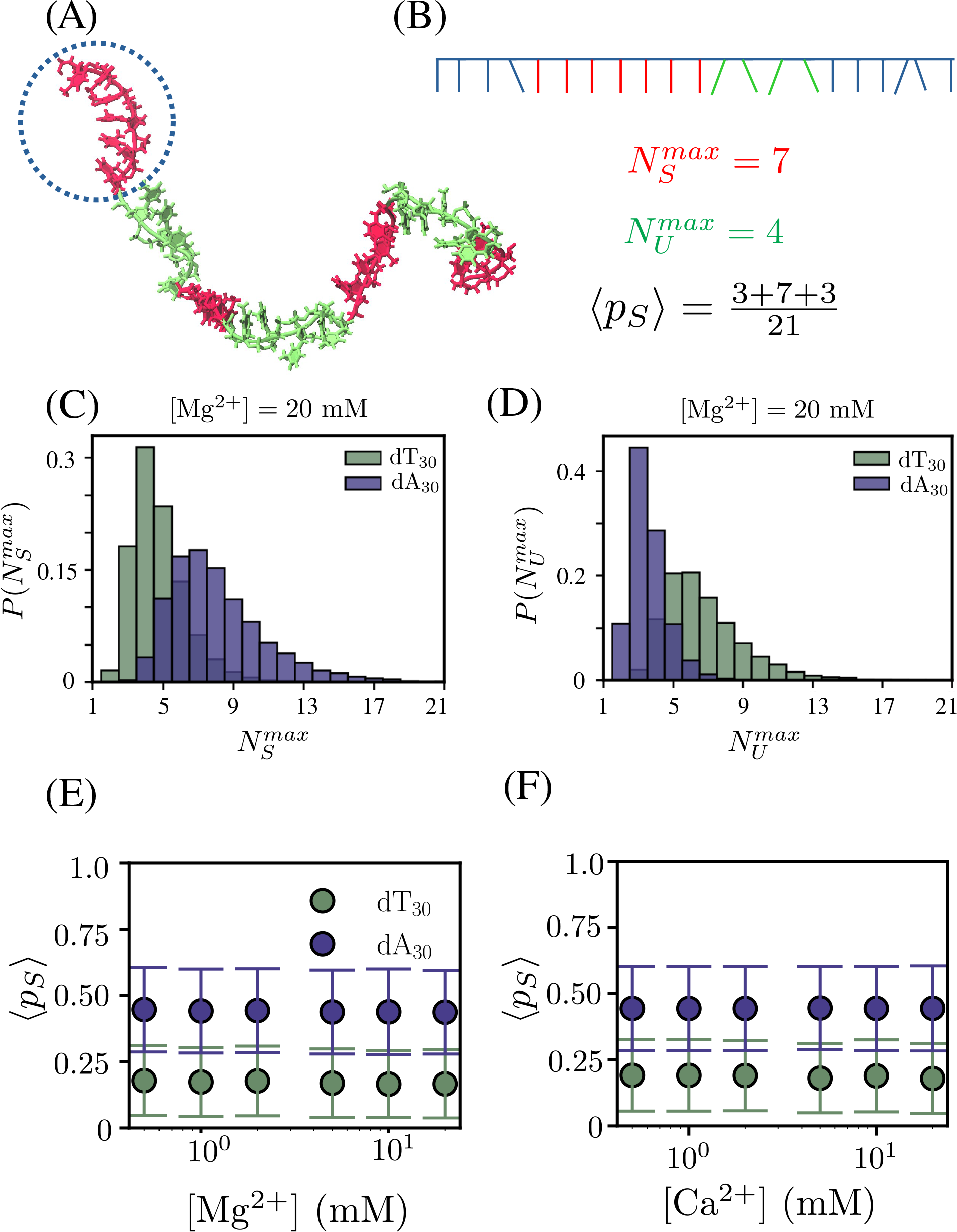
Stacking propensities in ssDNAs: (A) A simulation snapshot of dT_30_ chain highlighting a region of persistent stacks composed of six nucleotides. Stacks are shown in red. Unstacked regions are in green. (B) Schematic of a ssDNA segment depicting a stretch of the longest stacked bases, 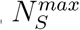(in red), stretch of longest unstacked bases, 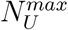 (in green), and ⟨*p*_*S*_⟩. (C) Normalized distributions of 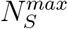 in dT_30_ and dA_30_ at [Mg^2+^] = 20 mM. (D) Same as (C), except the distributions correspond to 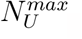. (E) Ensemble averaged values of the fraction of persistent stacks, ⟨*p*_*S*_⟩, as a function of [Mg^2+^]. (F) Same as (E), except the results are for [Ca^2+^].

### Spatial distribution of ions around dsDNA

The finding that 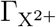 is the same for Mg^2+^ and Ca^2+^ (Figure 2) is surprising because one would expect that the number of smaller Mg^2+^ around the dsDNA should be larger than Ca^2+^. Hence, it is natural to ask if there are any discernible differences in the distributions of Mg^2+^ and Ca^2+^ ions around stiff dsDNAs at the nucleotide level which are masked in ion-counting and SAXS experiments? To answer this question we projected the ensemble averaged occupancy of ions onto the frontal plane (see Figure 9A) passing through the dsDNA at [X^2+^] = 20 mM. The two-dimensional projection of Mg^2+^ occupancy (Figure 9B) shows the cylindrical arrangement of Mg^2+^ ions in multiple layers surrounding the dsDNA. The inner-most layer contains a nearly uniform density of ions at a distance ≈ 0.85 nm from the center line of dsDNA, indicating non-specific binding, while ions are excluded from the backbone due to the finite sizes of DNA and ions. Unlike Mg^2+^, a relatively sparse layer of Ca^2+^ ions is localized at ≈ 0.95 nm from the center of line of dsDNA. In addition, Ca^2+^ ions are excluded to a greater extent from the dsDNA backbone compared to Mg^2+^ because of their larger size. Local ion concentrations 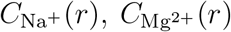 and 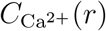, calculated using Eq. S1, are shown in the Figure S9 in the SI, also recapitulates these differences in ion binding. In agreement with the ion occupancy results, the highest [Mg^2+^] in the vicinity of the dsDNA is 520 mM at *r* ≈ 0.85 nm, whereas the highest [Ca^2+^] ≈ 450 mM at r ≈ 0.95 nm.

**Figure 9:**
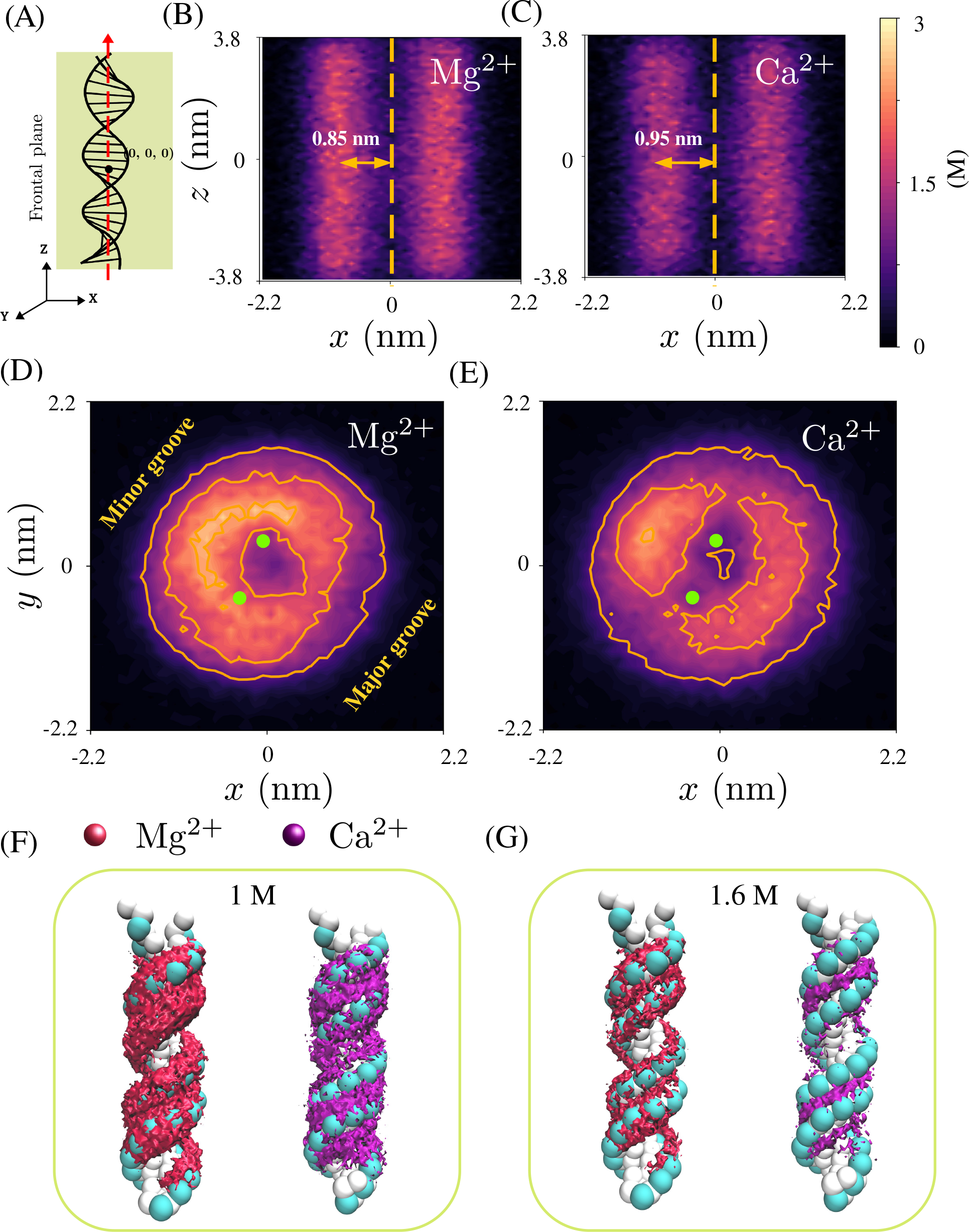
Two-dimensional spatial distribution of divalent cations around ds-DNA. (A) Illustration of the frontal plane passing through the dsDNA onto which ion occupancy is projected. Frontal plane is projected along the major principal axis, z, and the minor principal axis, *x*. The major principal axis is shown in red dashed line, passing through the origin. (B) Spatial distribution profile for Mg^2+^ ions projected onto the frontal plane passing through the dsDNA. (C) Same as (B), except the results are for Ca^2+^ ions. The local concentration scales are shown on the right. (D) 2D untwisted density of Mg^2+^ ions projected onto an average base-pair plane obtained by removing the helical twist [12], in [Mg^2+^] = 20 mM. Averaged locations of the phosphate groups are shown in green. Major and minor grooves are highlighted. (E) Same as (D), except results are shown for Ca^2+^ ions.(F)Ensemble average occupancy of Mg^2+^ (in red) and Ca^2+^ (in magenta) around dsDNA, in [X^2+^] = 20 mM. Isosurfaces are for grids with *ρ*_*i*_ = 1 M. Phosphate beads are highlighted in cyan. (G) Same as (F), except isosurfaces are for grids with *ρ*_*i*_ = 1.6 M.

The differences in spatial binding of Mg^2+^ and Ca^2+^ with phosphates, may be rationalized using the size differences alone. The size of Ca^2+^ ion (*R* ≈ 0.17 nm) is larger than Mg^2+^ ions (R ≈ 0.08 nm). As a result, the volume excluded by Ca^2+^ is eight times larger than Mg^2+^. The larger size also impacts the surface charge density of Ca^2+^, which is 4.5 times smaller than Mg^2+^, resulting in considerably weaker binding with the phosphate groups. Unlike the divalent cations, which show ≈ 20 fold increase in local concentration around dsDNA compared to bulk, maximum [Na^+^] ≈ 45 mM in Mg^2+^ solution, and ≈ 55 mM in Ca^2+^ solution at *r* ≈ 1.15 nm. This is a consequence of the lower charge density of the monovalent cations.

### Binding patterns of Mg^2+^ and Ca^2+^ to the grooves of dsDNA

To discern the groove specific binding of the divalent cations, the density of X^2+^ ions is projected onto *xy* plane of a base-pair. This is achieved by removing the helical twist angle for each base-pair plane, that is perpendicular to the helical axis (*z*), following a strategy discussed previously [12]. Such a representation successfully differentiates between the binding of the ions to the major and minor grooves. The resulting 2D densities of Mg^2+^ and Ca^2+^ ions are shown in Figures 9D and 9E, respectively, in [X^2+^] = 20 mM. Figure 9D vividly shows that Mg^2+^ binds to both the minor groove and the phosphates, while Ca^2+^ binds specifically to the minor groove. This binding pattern remains unchanged at all other [X^2+^] (Figure S10).

We also calculated the spatial occupancy of Mg^2+^ and Ca^2+^ ions surrounding DNA using three-dimensional (3D) cubic grid of length 1 Å. The number density corresponding to a grid *i* is computed as, 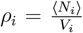, where ⟨*N*_*i*_⟩ is the average number of X^2+^ ions occupying the *i*^th^ grid with volume *V*_*i*_. Isosurfaces are drawn for Mg^2+^ (in red) and Ca^2+^ (in magenta) ions. Figure 9F and 9G show two isosurfaces highlighting grids (*i*) with *ρ*_i_ = 1 M and 1.6 M, respectively. At *ρ*_*i*_ = 1 M (Figure 9F), binding of Mg^2+^ and Ca^2+^ is almost uniform along the dsDNA cylinder. In contrast, at *ρ*_*i*_ = 1.6 M, Ca^2+^ ions are preferentially localized near the minor groove of DNA, whereas Mg^2+^ ions bind directly with the phosphate groups (Figure 9G), in addition to the minor grooves.

The binding modes are determined by the excluded volume of the divalent cations. The distance between adjacent phosphate groups belonging to the same strand is ≈ 0.6 nm. Due to smaller size of Mg^2+^ ions, it can directly interact with two phosphate groups (-1 e charge each), unlike Ca^2+^ ions. On the other hand, Ca^2+^ ions binding to the minor groove is dictated by the local phosphate ion concentration and the groove dimensions. In a typical B-DNA, the width of minor and major grooves are ≈ 1.2 nm and ≈ 2.2 nm, respectively. Due to close proximity of phosphate groups from two strands in the minor grooves, negative charge density is higher in the minor grooves compared to the major grooves. In addition, the width of the minor groove (≈ 1.2 nm) is sufficient to accommodate multiple Ca^2+^ ions which can neutralize phosphates from complementary strands simultaneously (Figure 9G).

### Spatial distribution of ions around ssDNA

To elucidate the spatial organization of ions along the ssDNA contour, we calculated a joint distribution function of two order parameters, *r*_*z*_ and 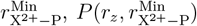, where 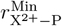 is the distance of closest approach between X^2+^ ions and the phosphate beads, and *r*_*z*_ is the *z*-component of the distance of the X^2+^ ion from the center of the major principal axis (presumed to lie along the *z* axis) of the ssDNA (see Figure 10A). To compare the spatial arrangement of Mg^2+^ ions around dT_30_ and dA_30_, we computed the difference in the joint distribution functions, 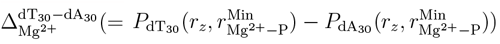 at 20 mM Mg^2+^ (Figure 10B). As is evident, Mg^2+^ ions preferentially accumulate along the contour of dT_30_ as compared to dA_30_. Lack of persistent structural order increases the local phosphate charge density in dT_30_, as opposed to dA_30_, which is rich in helical motifs (Figures 6, 7). A similar trend is observed for Ca^2+^ ions as well (Figure 10C), except that the highest Mg^2+^ occupancy is found at 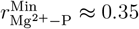 nm, whereas for Ca^2+^, it is located at 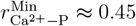 nm, due to its larger size.

**Figure 10:**
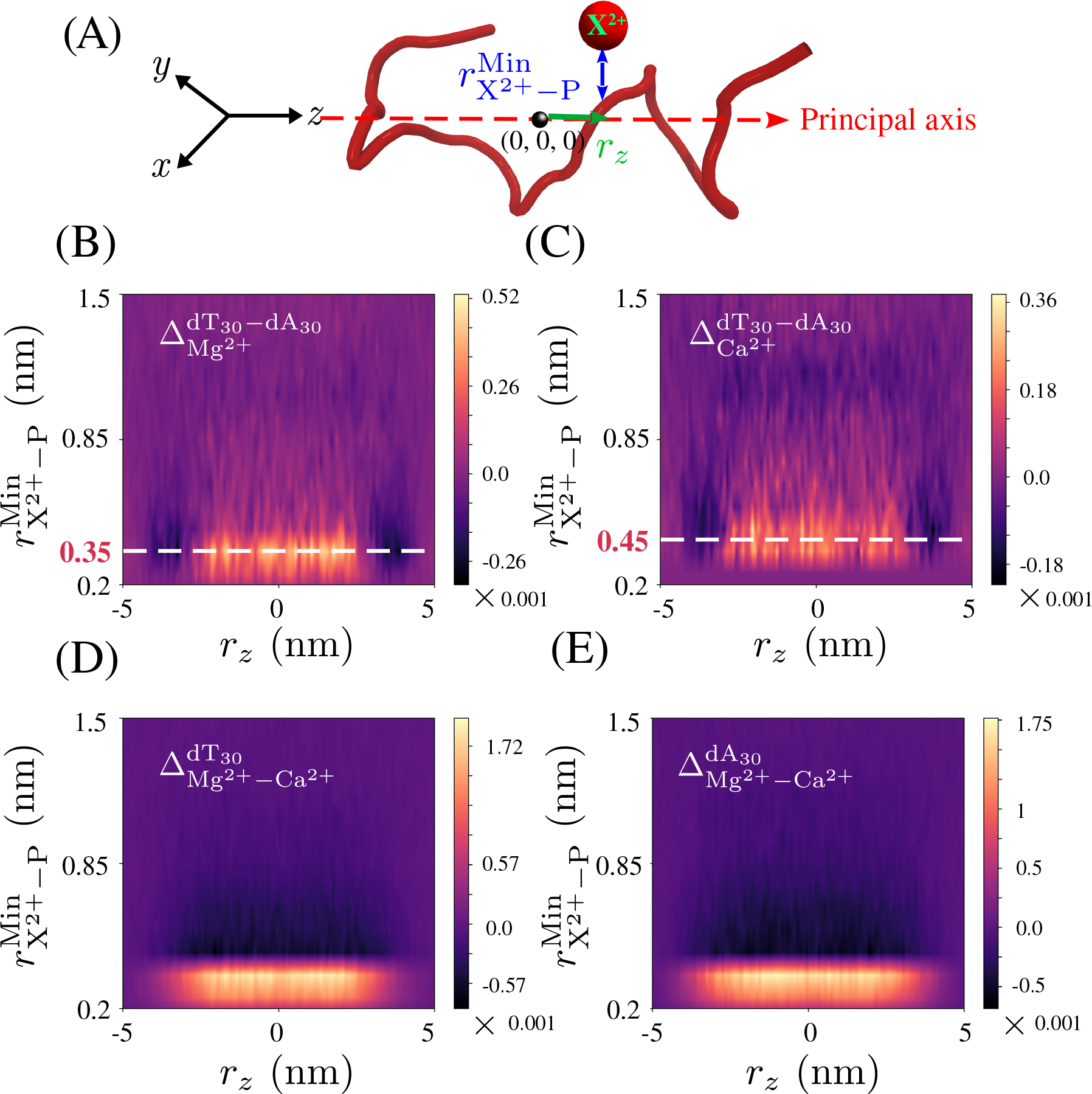
Two-dimensional spatial distribution of divalent cations around ssDNA. (A) Definition of 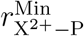 and *r*_*z*_, using a typical ssDNA construct. Major principal axis (red dashed line) is aligned along the *z* axis. Ion X^2+^ is in red, 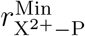 is the distance of closest approach between the phosphate and X^2+^, and *r*_*z*_ is the *z*-component of distance of X^2+^ from the center of the major principal axis of ssDNA. (B) Difference in the normalized joint distribution functions, 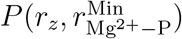, for dT_30_ and dA_30_, 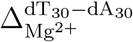, in [Mg^2+^] = 20 mM. (C) Same as (B), except the results correspond to are Ca^2+^ ions. (D) Difference in 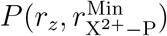, for dT_30_, in [Mg^2+^] = 20 mM and [Ca^2+^] = 20 mM, 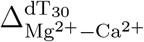. (E) Same as (D), except the plot is for dA_30_.

We compared the relative population of Mg^2+^ and Ca^2+^ ions, around dT_30_, by computing the difference in the joint distribution functions, 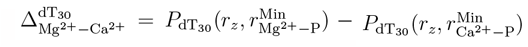 (Figure 10D). The population of Ca^2+^ ions is reduced in the immediate vicinity of dT_30_, compared to Mg^2+^, indicating weaker interactions and more diffuse binding. Figure 10E illustrates the results for dA_30_, where qualitatively similar differences are found between Mg^2+^ and Ca^2+^ ions.

To quantify the local ion condensation, we calculated the concentration of ion X^2+^ in the vicinity of individual phosphate groups *i*, 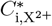, using Eq. S3 in the SI, within a spherical volume of radius set by the Bjerrum length *l*_*B*_ (Figure S11). For 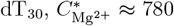 mM, is larger than 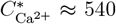 mM, which is a consequence of higher charge density and smaller size of Mg^2+^. A similar trend is found in dA_30_, where 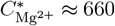 mM, whereas 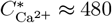 mM. In both the cases, there are non-specific interactions between the phosphate groups and X^2+^ ions, which may be attributed to the lack of tertiary structures in the homopolymeric sequences. This non-specificity is evident when the location of X^2+^ is analyzed around each nucleotide (direct interactions only) in individual conformations (Figures S13, S14).

## DISCUSSION

### Mg^2+^ and Ca^2+^ sense major and minor grooves in dsDNA differently

Neither the dependence of 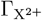 on [X^2+^] (Figures 2 and 3) nor the scattering profiles (Figures 4 and 5) show significant differences between [Mg^2+^] and [Ca^2+^]. Therefore, the conformations of the short dsDNA sequences are determined predominantly by the stacking and base-pairing interactions, with electrostatic interactions playing a sub-dominant role. Divalent cations merely decrease the effective charge on the phosphate groups. Thus, 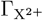 and I(q) are impervious to the nature of the divalent cations. However, differences between [Mg^2+^] and [Ca^2+^] emerge when the details of their binding to dsDNA are examined at a finer scale: [Mg^2+^] binds predominantly to the phosphate groups and minor grooves, whereas [Ca^2+^] preferentially binds to the minor groove, and exhibits little or no direct interactions with the phosphate groups. The distinct binding modes of Mg^2+^ and Ca^2+^ to dsDNA could affect the differential interaction of proteins with DNA [60], and possibly certain enzymatic activities where Ca^2+^ can not substitute for Mg^2+^ [61, 62].

### Stacking versus electrostatic interactions in ssDNA

The contrast in the extent of heterogeneity between dT_30_ and dA_30_ is striking. The stability of ssDNA should result due to the interplay between favorable stacking interactions (U_ST_) and unfavorable electrostatic interactions (*U*_*EL*_). In dT_30_, these two interactions nearly cancel each other (Figure S7). As a result, dT_30_ behaves like a random coil, especially at high [X^2+^] (Figures 7E, F show *v*_*eff*_ ≈ 0.6 in 20 mM Mg^2+^ or Ca^2+^). In contrast, the stability of dA_30_ is determined by the favorable stacking interactions, which are strong enough to overcome the electrostatic repulsion (figure S7). As a result, dA_30_ forms helical structures at both [X^2+^] = 0 mM and 20 mM (Figures 7E, F).

Although we have focused on dT_30_ and dA_30_, our result could be applicable to dC_n_ and dG_*n*_. We will assume that n is not too large because as n increases there is a high probability that the can fold on itself, stabilized by the formation of intra-molecular stacks. It is thought (see for example [63]) that the order of stacking interactions, from the most to the favorable may be arranged as 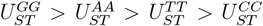. It follows from our results that dG_30_ is likely to form more stable and stiff helix over the range of ion concentrations examined here, whereas the conformations of dC_30_ would be determined almost exclusively by *U*_*EL*_. In other words, dC_30_ is, for practical purposes, should be viewed as a short polyelectrolyte.

## CONCLUSIONS

Using a transferable computational TIS-ION model that includes monovalent and divalent cations explicitly, we probed the interplay between stacking, base-pairing (dsDNA), and electrostatic interactions in modulating the conformations of dsDNA and ssDNA sequences. The energy function combines the coarse-grained TIS model for nucleic acids with explicit models for the spherical cations (Na^+^, Mg^2+^, and Ca^2+^). Calculations of the number of monovalent and divalent cations in the vicinity of dsDNA and ssDNA sequences, over a wide range of cation concentrations, are in excellent agreement with experiments [6, 10] as are the SAXS profiles for dsDNA, dT_30_ and dA_30_. Remarkably, the accurate predictions were made *without fitting* a single parameter to experimental data, thus attesting to the robustness and transferability of the model.

Strikingly, we find that neither SAXS nor ion-counting experiments distinguishes between Mg^2+^ and Ca^2+^, in terms of phosphate binding, but the fingerprints of preferential binding patterns are discernible at the molecular scale. While Mg^2+^ binds to both the phosphate groups, and DNA minor groove, the larger sized Ca^2+^ are exclusively localized near the minor groove. The interplay between stacking and electrostatic interactions is most evident in the nature of conformations explored by the ssDNAs, dT_30_ and dA_30_. In the former, these two interactions nearly cancel each other, which makes dT_30_ behave as a homopolymer in a good solvent, at sufficiently high divalent ion concentrations. On the other hand, because stacking dominates in dA_30_, the conformations have considerable helical order.

## Data Availability

Simulations were performed using an in-house code built on the OpenMM platform. The code is available at https://github.com/balaka92/TIS-DNA-Ex-Ion.

## Acknowledgements

This work was supported by the National Science Foundation through CHE 232056. Additional support from the Welch Foundation (F-0019) through the Collie-Welch Chair is gratefully acknowledged. We thank the Texas Advanced Computing Center and Pittsburgh Supercomputing Center for providing generous computational resources.

## References

[1] I. Rouzina and V. A. Bloomfield, DNA bending by small, mobile multivalent cations, Biophys. J. 74, 3152–3164 (1998).

[2] P. Soultanas, M. S. Dillingham, S. S. Velankar, and D. B. Wigley, DNA binding mediates conformational changes and metal ion coordination in the active site of PcrA helicase, J. Mol. Biol. 290, 137–148 (1999).

[3] S. A. Grigoryev, G. Arya, S. Correll, C. L. Woodcock, and T. Schlick, Evidence for heteromorphic chromatin fibers from analysis of nucleosome interactions, Proc. Natl. Acad. Sci. USA 106, 13317–13322 (2009).

[4] G. C. Wong and L. Pollack, Electrostatics of strongly charged biological polymers: Ion-mediated interactions and self-organization in nucleic acids and proteins, Annu. Rev. Phys. Chem. 61, 171–189 (2010).

[5] C. G. Baumann, S. B. Smith, V. A. Bloomfield, and C. Bustamante, Ionic effects on the elasticity of single DNA molecules, Proc. Natl. Acad. Sci. USA 94, 6185–6190 (1997).

[6] Y. Bai, M. Greenfeld, K. J. Travers, V. B. Chu, J. Lipfert, S. Doniach, and D. Herschlag, Quantitative and comprehensive decomposition of the ion atmosphere around nucleic acids, J. Am. Chem. Soc. 129, 14981–14988 (2007).

[7] S. P. Meisburger, J. L. Sutton, H. Chen, S. A. Pabit, S. Kirmizialtin, R. Elber, and L. Pollack, Polyelectrolyte properties of single stranded DNA measured using SAXS and single-molecule FRET: Beyond the wormlike chain model, Biopolymers 99, 1032–1045 (2013).

[8] M. C. Murphy, I. Rasnik, W. Cheng, T. M. Lohman, and T. Ha, Probing single-stranded DNA conformational flexibility using fluorescence spectroscopy, Biophys. J. 86, 2530–2537 (2004).

[9] D. B. McIntosh, G. Duggan, Q. Gouil, and O. A. Saleh, Sequence-dependent elasticity and electrostatics of single-stranded DNA: Signatures of base-stacking, Biophys. J. 106, 659–666 (2014).

[10] A. Plumridge, S. P. Meisburger, K. Andresen, and L. Pollack, The impact of base stacking on the conformations and electrostatics of single-stranded DNA, Nucleic Acids Res. 45, 3932–3943 (2017).

[11] P. Yakovchuk, E. Protozanova, and M. D. Frank-Kamenetskii, Base-stacking and base-pairing contributions into thermal stability of the DNA double helix, Nucleic Acids Res. 34, 564–574 (2006).

[12] G. M. Giambasu, T. Luchko, D. Herschlag, D. M. York, and D. A. Case, Ion counting from explicit-solvent simulations and 3D-RISM, Biophys. J. 106, 883–894 (2014).

[13] A. Plumridge, S. P. Meisburger, and L. Pollack, Visualizing single-stranded nucleic acids in solution, Nucleic Acids Res. 45, e66 (2017).

[14] O. A. Saleh, D. B. McIntosh, P. Pincus, and N. Ribeck, Nonlinear low-force elasticity of singlestranded DNA molecules, Phys. Rev. Lett. 102, 068301 (2009).

[15] N. M. Toan and D. Thirumalai, Theory of biopolymer stretching at high forces, Macromolecules 43, 4394–4400 (2010).

[16] I. A. Shkel, O. V. Tsodikov, and M. T. Record Jr, Asymptotic solution of the cylindrical nonlinear Poisson–Boltzmann equation at low salt concentration: Analytic expressions for surface potential and preferential interaction coefficient, Proc. Natl. Acad. Sci. USA 99, 2597–2602 (2002).

[17] S. Kirmizialtin, A. R. Silalahi, R. Elber, and M. O. Fenley, The ionic atmosphere around A-RNA: Poisson-Boltzmann and molecular dynamics simulations, Biophys. J. 102, 829–838 (2012).

[18] G. Lamm and G. R. Pack, Local dielectric constants and Poisson–Boltzmann calculations of DNA counterion distributions, Int. J. Quantum Chem. 65, 1087–1093 (1997).

[19] V. B. Chu, Y. Bai, J. Lipfert, D. Herschlag, and S. Doniach, Evaluation of ion binding to DNA duplexes using a size-modified Poisson-Boltzmann theory, Biophys. J. 93, 3202–3209 (2007).

[20] A. Vologodskii and N. Cozzarelli, Modeling of long-range electrostatic interactions in DNA, Biopolymers 35, 289–296 (1995).

[21] N. A. Denesyuk and D. Thirumalai, Coarse-grained model for predicting RNA folding thermodynamics., J. Phys. Chem. B 117, 4901–4911 (2013).

[22] D. Chakraborty, N. Hori, and D. Thirumalai, Sequence-dependent three interaction site model for single-and double-stranded DNA, J. Chem. Theory Comput. 14, 3763–3779 (2018).

[23] H. T. Nguyen, N. Hori, and D. Thirumalai, Theory and simulations for RNA folding in mixtures of monovalent and divalent cations, Proc. Natl. Acad. Sci. USA 116, 21022–21030 (2019).

[24] L. Z. Sun, J. L. Qian, P. Cai, and X. Xu, Mutual effects between single-stranded DNA conformation and Na+-Mg2+ ion competition in mixed salt solutions, Phys. Chem. Chem. Phys. 24, 20867–20881 (2022).

[25] A. Savelyev and A. D. MacKerell Jr, Competition among Li^+^, Na^+^, K^+^, and Rb^+^ monovalent ions for DNA in molecular dynamics simulations using the additive CHARMM36 and drude polarizable force fields, J. Phys. Chem. B 119, 4428–4440 (2015).

[26] L. A. Yates, R. J. Aramayo, N. Pokhrel, C. C. Caldwell, J. A. Kaplan, R. L. Perera, M. Spies, E. Antony, and X. Zhang, A structural and dynamic model for the assembly of Replication Protein A on single-stranded DNA, Nat. Commun. 9 (2018).

[27] S. Cruz-León, K. K. Grotz, and N. Schwierz, Extended magnesium and calcium force field parameters for accurate ion–nucleic acid interactions in biomolecular simulations, J. Chem. Phys. 154, 171102 (2021).

[28] J. Yoo and A. Aksimentiev, Competitive binding of cations to duplex DNA revealed through molecular dynamics simulations, J. Phys. Chem. B 116, 12946–12954 (2012).

[29] D. M. Hinckley and J. de Pablo, Coarse-grained ions for nucleic acid modeling, J. Chem. Theory Comput. 11, 5436–5446 (2015).

[30] C. Hyeon and D. Thirumalai, Mechanical unfolding of RNA hairpins, Proc. Natl. Acad. Sci. USA 102, 6789–6794 (2005).

[31] C. Hyeon and D. Thirumalai, Capturing the essence of folding and functions of biomolecules using coarse-grained models, Nat. Commun. 2, 487 (2011).

[32] T. E. Ouldridge, A. A. Louis, and J. P. Doye, Structural, mechanical, and thermodynamic properties of a coarse-grained DNA model, J. Chem. Phys. 134, 085101 (2011).

[33] D. M. Hinckley, G. S. Freeman, J. K. Whitmer, and J. de Pablo, An experimentally-informed coarse-grained 3-site-per-nucleotide model of DNA: Structure, thermodynamics, and dynamics of hybridization, J. Chem. Phys. 139, 144903 (2013).

[34] L. Z. Sun, J. L. Qian, P. Cai, H. X. Hu, X. Xu, and M. B. Luo, Mg^2+^ effects on the singlestranded DNA conformations and nanopore translocation dynamics, Polymer 250, 124895 (2022).

[35] N. A. Denesyuk and D. Thirumalai, How do metal ions direct ribozyme folding?, NatChem 7, 793–801 (2015).

[36] N. Hori, N. A. Denesyuk, and D. Thirumalai, Shape changes and cooperativity in the folding of the central domain of the 16S ribosomal RNA, Proc. Natl. Acad. Sci. USA 118, e2020837118 (2021).

[37] N. Hori and D. Thirumalai, Watching ion-driven kinetics of ribozyme folding and misfolding caused by energetic and topological frustration one molecule at a time, Nucleic Acids Res. 51, gkad755 (2023).

[38] J. SantaLucia, H. T. Allawi, and P. A. Seneviratne, Improved nearest-neighbor parameters for predicting DNA duplex stability, Biochemistry 35, 3555–3562 (1996).

[39] J. SantaLucia and D. Hicks, The thermodynamics of DNA structural motifs, Annu. Rev. Biophys. Biomol. Struct. 33, 415–440 (2004).

[40] J. Hasted, Water: A comprehensive treatise, The Physics and Physical Chemistry of Water 1, 255–305 (1972).

[41] P. Eastman, J. Swails, J. D. Chodera, R. T. McGibbon, Y. Zhao, K. A. Beauchamp, L. P. Wang, A. C. Simmonett, M. P. Harrigan, C. D. Stern, R. P. Wiewiora, B. R. Brooks, and V. S. Pande, OpenMM 7: Rapid development of high performance algorithms for molecular dynamics, PLOS Biol. 13, e1005659 (2017).

[42] T. Darden, D. York, and L. Pedersen, Particle mesh Ewald: An Nlog(N) method for Ewald sums in large systems, J. Chem. Phys. 98, 10089–10092 (1993).

[43] J. D. Honeycutt and D. Thirumalai, The nature of folded states of globular proteins, Biopolymers 32, 695–709 (1992).

[44] R. W. Hockney and J. W. Eastwood, Computer simulation using particles (CRC Press, 2021).

[45] S. A. Pabit, X. Qiu, J. S. Lamb, L. Li, S. P. Meisburger, and L. Pollack, Both helix topology and counterion distribution contribute to the more effective charge screening in dsRNA compared with dsDNA, Nucleic Acids Res. 37, 3887–3896 (2009).

[46] P. Smith, Equilibrium dialysis data and the relationships between preferential interaction parameters for biological systems in terms of Kirkwood-Buff integrals, J. Phys. Chem. B 110, 2862–2868 (2006).

[47] J. Schurr, D. Rangel, and S. Aragon, A contribution to the theory of preferential interaction coefficients, Biophys. J. 89, 2258–2276 (2005).

[48] D. K. Klimov, M. R. Betancourt, and D. Thirumalai, Virtual atom representation of hydrogen bonds in minimal off-lattice models of alpha helices: Effect on stability, cooperativity and kinetics, Folding and Design 3, 481–496 (1998).

[49] V. A. Bloomfield, Condensation of DNA by multivalent cations: Considerations on mechanism, Biopolymers 31, 1471–1481 (1991).

[50] P. Arscott, C. MA J. Wenner, and V. Bloomfield, DNA condensation by cobalt hexaammine(III) in alcohol-water mixtures - dielectric-constant and other solvent effects, Biopolymers 36, 345–364 (1995).

[51] C. Ma and V. A. Bloomfield, Gel electrophoresis measurement of counterion condensation on DNA, Biopolymers 35, 211–216 (1995).

[52] R. W. Wilson and V. A. Bloomfield, Counterion-induced condensation of deoxyribonucleic acid. A light-scattering study, Biochemistry 18, 2192–2196 (1979).

[53] V. A. Bloomfield, DNA condensation by multivalent cations, Biopolymers 44, 269–282 (1997).

[54] A. Rohatgi, Webplotdigitizer: Version 4.6 (2022).

[55] N. Hori, TIS2AA (2017).

[56] D. Svergun, C. Barberato, and M. H. Koch, CRYSOL - A program to evaluate X-ray solution scattering of biological macromolecules from atomic coordinates, J. Appl. Crystallogr. 28, 768–773 (1995).

[57] J. S. Higgins and H. C. Benoit, Polymers and neutron scattering (Oxford University Press, London, 1994).

[58] A. Y. Sim, J. Lipfert, D. Herschlag, and S. Doniach, Salt dependence of the radius of gyration and flexibility of single stranded DNA in solution probed by Small-Angle X-ray Scattering, Phys. Rev. E 86, 021901 (2012).

[59] C. Ke, M. Humeniuk, S. Hanna, and P. E. Marszalek, Direct measurements of base stacking interactions in DNA by single-molecule atomic-force spectroscopy, Phys. Rev. Lett. 99, 018302 (2007).

[60] M. Osawa, A. Dace, K. I. Tong, A. Valiveti, M. Ikura, and J. B. Ames, Mg^2+^ and Ca^2+^ differentially regulate DNA binding and dimerization of DREAM, J. Biol. Chem. 280, 18008–18014 (2005).

[61] S. R. Bellamy, Y. S. Kovacheva, I. H. Zulkipli, and S. E. Halford, Differences between Ca2+ and Mg2+ in DNA binding and release by the SfiI restriction endonuclease: implications for DNA looping, Nucleic Acids Res. 37, 5443–5453 (2009).

[62] A. Pingoud, M. Fuxreiter, V. Pingoud, and W. Wende, Type II restriction endonucleases: structure and mechanism, Cell. Mol. Life Sci. 62, 685–707 (2005).

[63] R. F. Brown, C. T. Andrews, and A. H. Elcock, Stacking free energies of all DNA and RNA nucleoside pairs and dinucleoside-monophosphates computed using recently revised AMBER parameters and compared with experiment, J. Chem. Theory Comput. 11, 2315–2328 (2015).

[64] T. J. Macke and D. A. Case, Modeling unusual nucleic acid structures, ACS Publications 682, 379–393 (1998).

[65] W. Humphrey, A. Dalke, and K. Schulten, VMD: Visual molecular dynamics, J. Mol. Graph. 14, 33–38 (1996).

