## Supplementary material for "Competition between stacking and divalent cation mediated electrostatic interactions determines the conformations of short DNA sequences"

**Calculation of local ion concentration profiles near DNA:** The rigid 24 base pair (bp) dsDNA has cylindrical symmetry around the principal $z$-axis that is aligned along the length of the cylinder. In contrast, the flexible ssDNA sequences is spherically symmetric around their centers of mass (see Figure S2). The local concentration $C_X(r)$ associated with ion $X$, as a function of distance $r$ from the DNA is computed using,

$$C_X(r) = \frac{N_X(r)}{\Delta V_r C_X^S} \tag{S1}$$

where $N_X(r)$ is the number of ion $X$ within a volume $\Delta V_r$ and $C_X^S$ is the bulk concentration. For dsDNA, $r$ is the radial distance of the ion from the principal axis ($z$), and $\Delta V_r$ is volume of the cylindrical shell between $r$ and $r + \Delta r$. Similarly, for ssDNA, $r$ is the distance of the ion from the center of mass, and $\Delta V_r$ is the volume of the spherical shell between $r$ and $r + \Delta r$.

**Effective bulk concentration $C^B$ :** Because DNA is highly charged, both monovalent and divalent cations are attracted towards the phosphate groups, thus causing a deficit of cations in the bulk. As a result, the effective bulk concentration, $C_X^B$, differs from the actual concentration $C_X^S$. The latter is obtained by dividing the total number of $X$ ions by the system volume. The plateau value in the concentration profile $C_X(r)$ is taken to be $C_X^B$,

$$C_X^B = \langle C_X(r) \rangle \text{ when } \frac{dC_X(r)}{dr} \approx 0. \tag{S2}$$

**Local concentration of ions around phosphate:**

We calculated the local ion concentration (in molar units) around a phosphate group of the $i^{th}$ nucleotide $C_{i,X}^*$, using,

$$C_{i,X}^* = \frac{1}{N_A V_c} \int_0^{r_c} \rho_{i,X}(r) 4\pi r^2 dr, \tag{S3}$$

where $\rho_{i,X}(r)$ is the number density of the ion at a distance $r$ from the phosphate bead, $V_c$ is the volume of a sphere of radius $r_c$, and $N_A$ is the Avogadro's number. We use the Bjerrum length (at $T = 277$ K) as the cutoff distance, $r_c = l_B = 0.698$ nm, where $l_B$ is the distance at which Coulomb attraction between a cation and an anion ($Z = 1$) is $k_B T$.

**Small Angle X-Ray Scattering (SAXS) profiles from coarse-grained trajectories:** The SAXS profiles were calculated by first converting ≈ 10,000 randomly chosen conformations generated in the coarse-grained simulations to all-atom details, using an in-house code TIS2AA[1] that uses a DNA-fragment library. Subsequently, the energy of the all atom structures were minimized using AMBER-99 bsc0[2] force field with OL15[3] corrections. Using the pool of all-atom structures, we calculated the SAXS profiles for dsDNA and ssDNA sequences as a function of [$Mg^{2+}$] or [$Ca^{2+}$].

We calculated the scattering intensity, $I(q)$, as a function of the momentum transfer vector $q$, from the atomic coordinates with $N$ heavy atoms using the Debye equation,

$$I(q) = \sum_{i=1}^{N} \sum_{j=1}^{N} f_i(q) f_j(q) \frac{\sin(q \cdot r_{ij})}{q \cdot r_{ij}}, \tag{S4}$$

where $f_i(q)$ and $f_j(q)$ are the form factors associated with the $i^{th}$ and $j^{th}$ atoms separated by a distance $r_{ij}$ at $q$. We used the software CRYSOL[4] to construct the SAXS profiles using the all-atom trajectories with default hydration parameters. In CRYSOL, spherically averaged form-factors are computed for each atomic group in the range $0 < q < 10$ nm$^{-1}$ using a five-Gaussian approximation of the form factors of individual atoms and inter-atomic distances. *International Tables for X-ray Crystallography* (1974) lists the form factors of the individual atoms.[5]

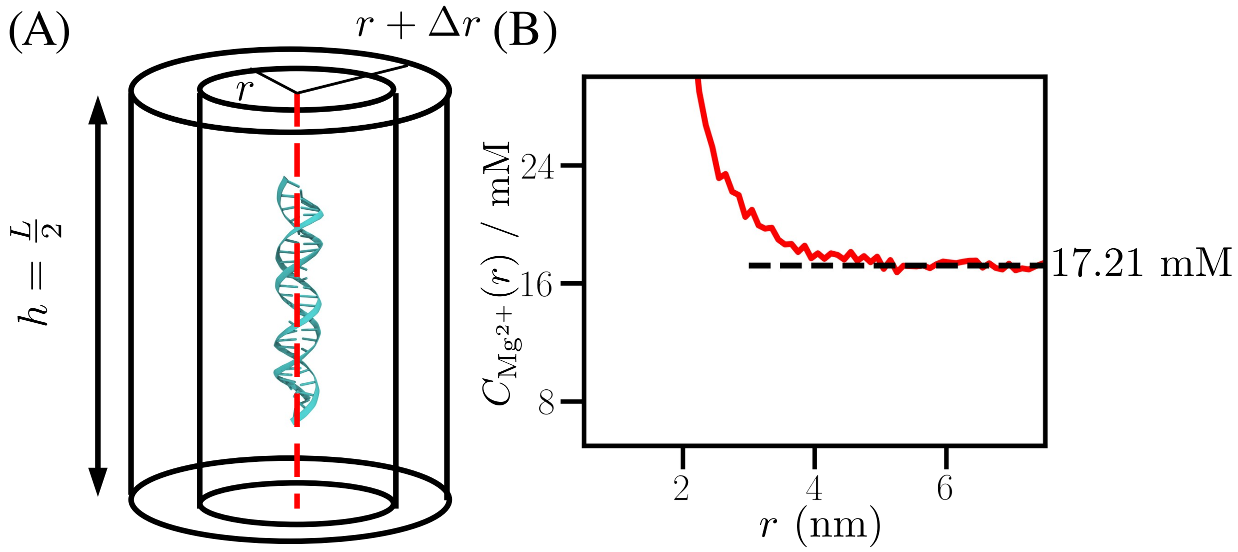


Figure S1: (A) Local concentration $C_X(r)$ for dsDNA calculated using Eq. S1 between $r$ and $r + \Delta r$. (B) Bulk concentration of $\mathrm{Mg^{2+}}$ ion ($C^B_{\mathrm{Mg^{2+}}}$), is calculated at large $r$ where $\frac{dC_{\mathrm{Mg^{2+}}}(r)}{dr} \approx 0$. In the example shown, $C^S_{\mathrm{Mg^{2+}}} = 20$ mM while $C^B_{\mathrm{Mg^{2+}}} \approx 17.21$ mM.

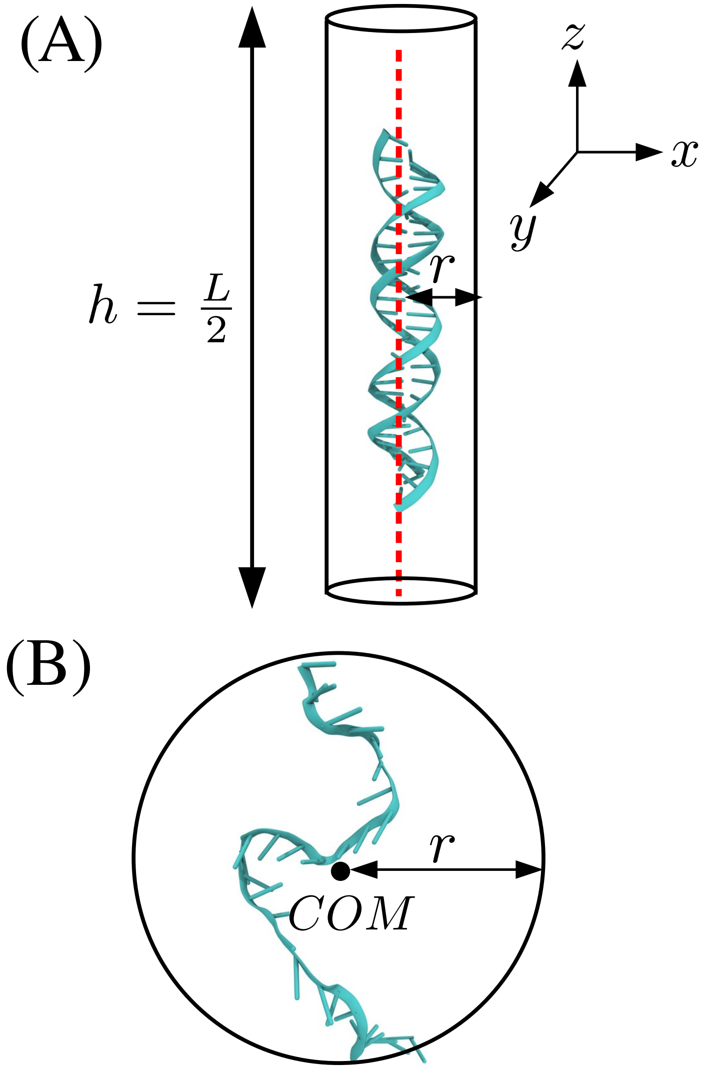


Figure S2: **Scheme for evaluating $\Gamma_i$ in Eq. 7 and 8 in the main text** (A) The radial distance, $r$, of an ion, is measured from the principal axis of dsDNA (red dashed line) where $h$ is the effective cylinder height. (B) For a ssDNA, $r$ is the distance of an ion from the center of mass of the NA.

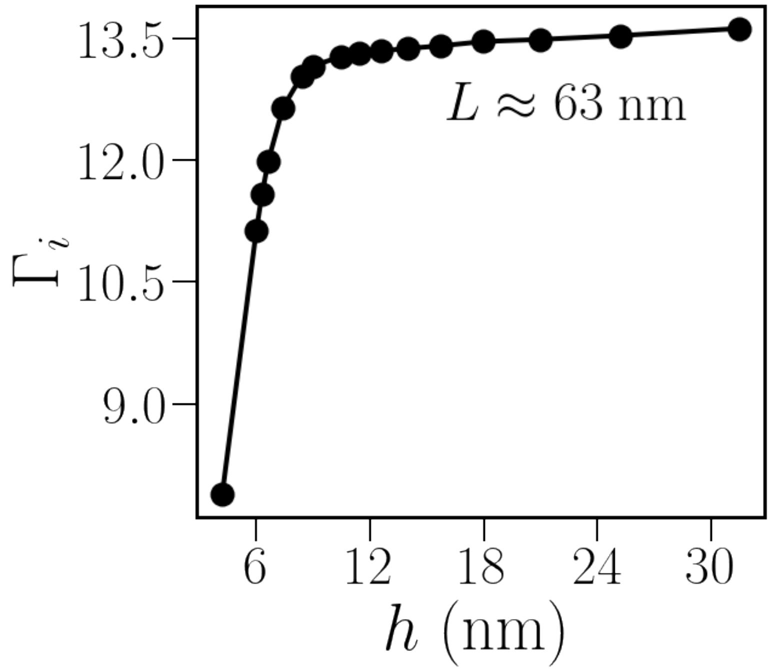


Figure S3: Variation of $\Gamma_i$ (see Eq. 7 in the main text) with height $h$ of the cylinder for dsDNA. Results shown for $[X^{2+}] = 0.1$ mM and box length $L = 63$ nm.

(A)

$[Mg^{2+}] = 3$ mM

(B)

$[Mg^{2+}] = 16$ mM

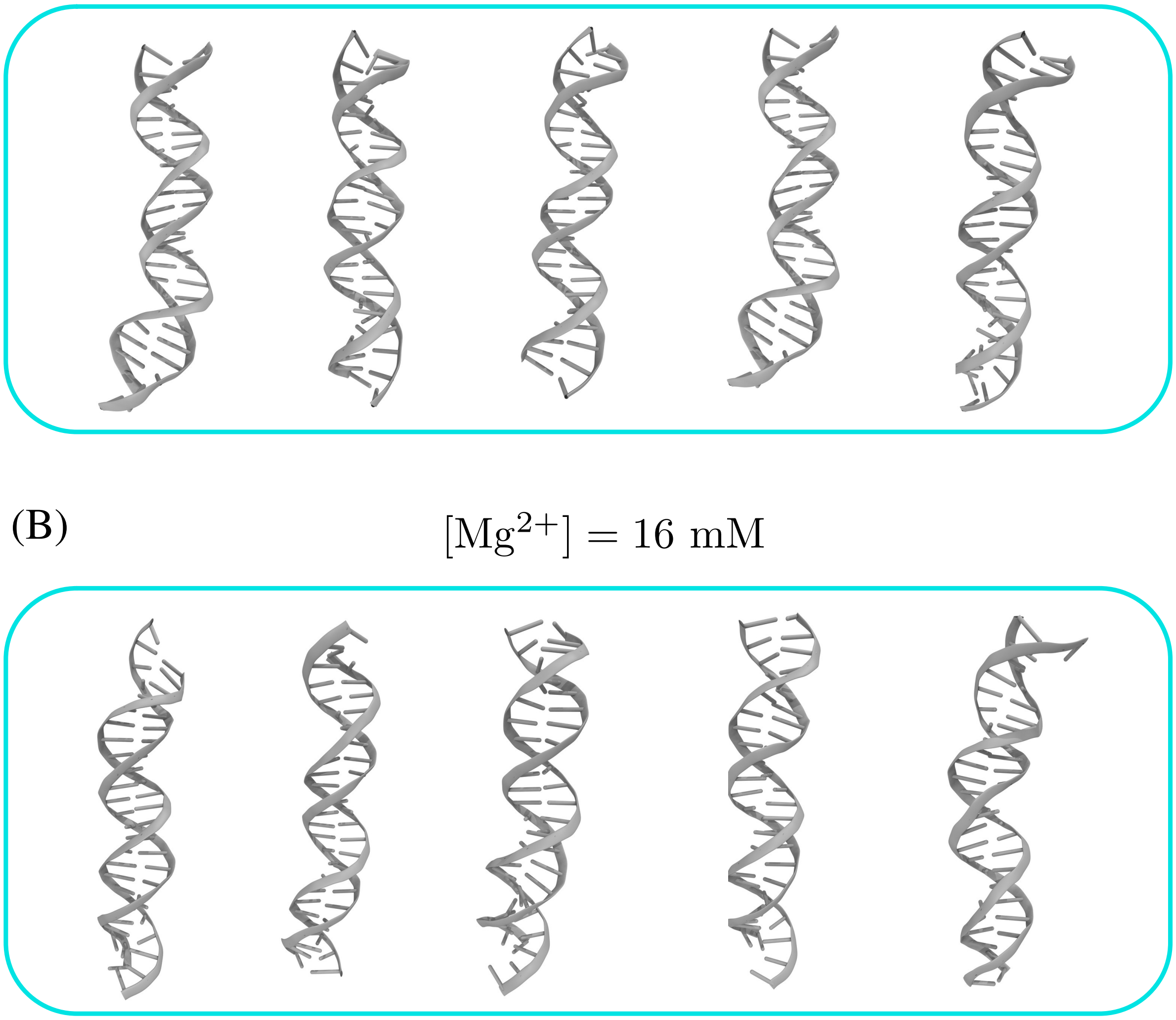

Figure S4: (A) Reconstructed all-atom structures from the coarse-graied simulations for the 25 bp dsDNA used to compute SAXS profiles in $[Mg^{2+}] = 3$ mM. (B) Same as (A), except for $[Mg^{2+}] = 16$ mM.

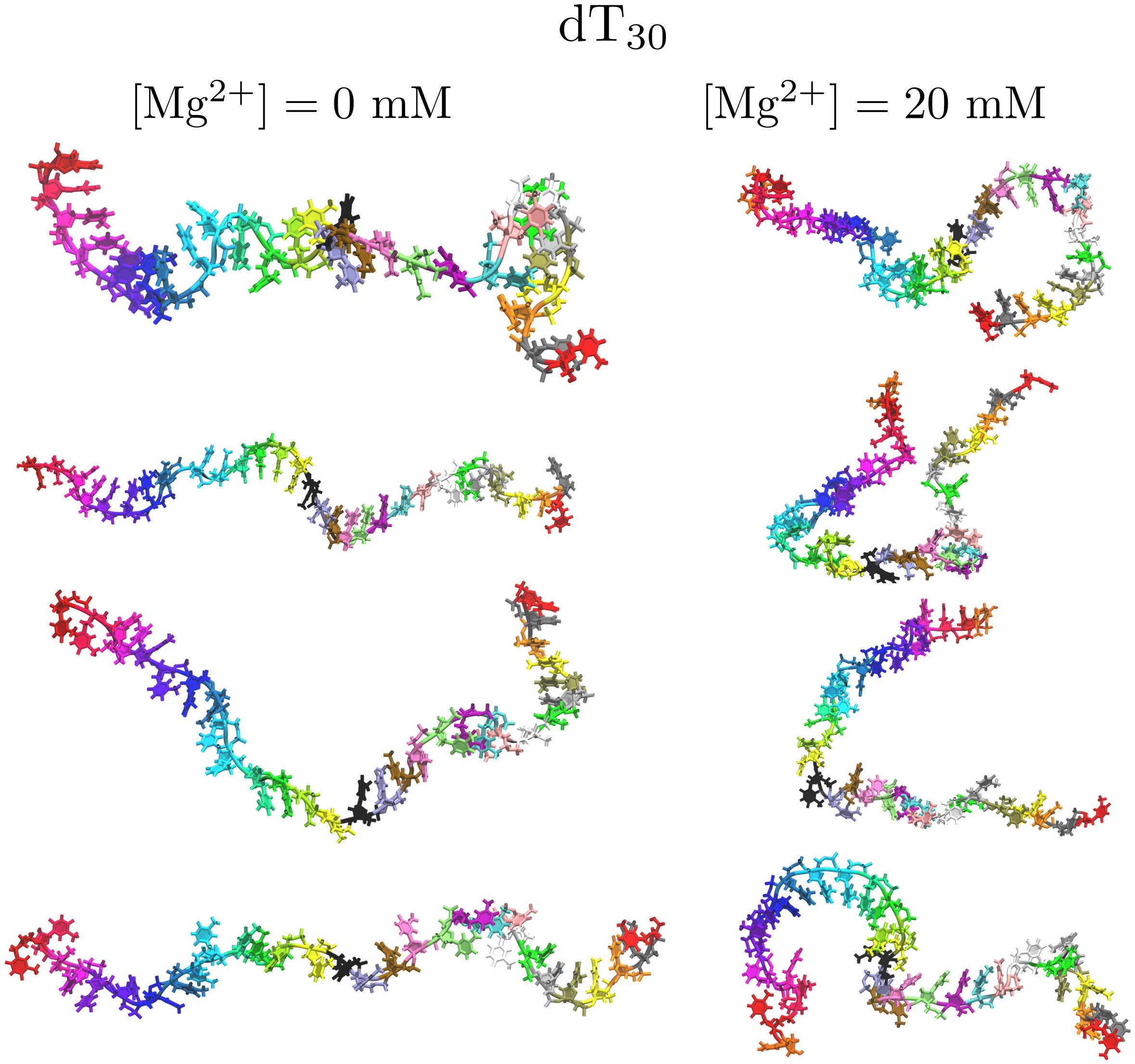


Figure S5: Reconstructed all-atom structures for $dT_{30}$ from coarse-grained simulations, used in computing SAXS profiles, in $[Mg^{2+}] = 0$ mM and 20 mM. To aid visualization, nucleotides are assigned different colors based on their sequence number.

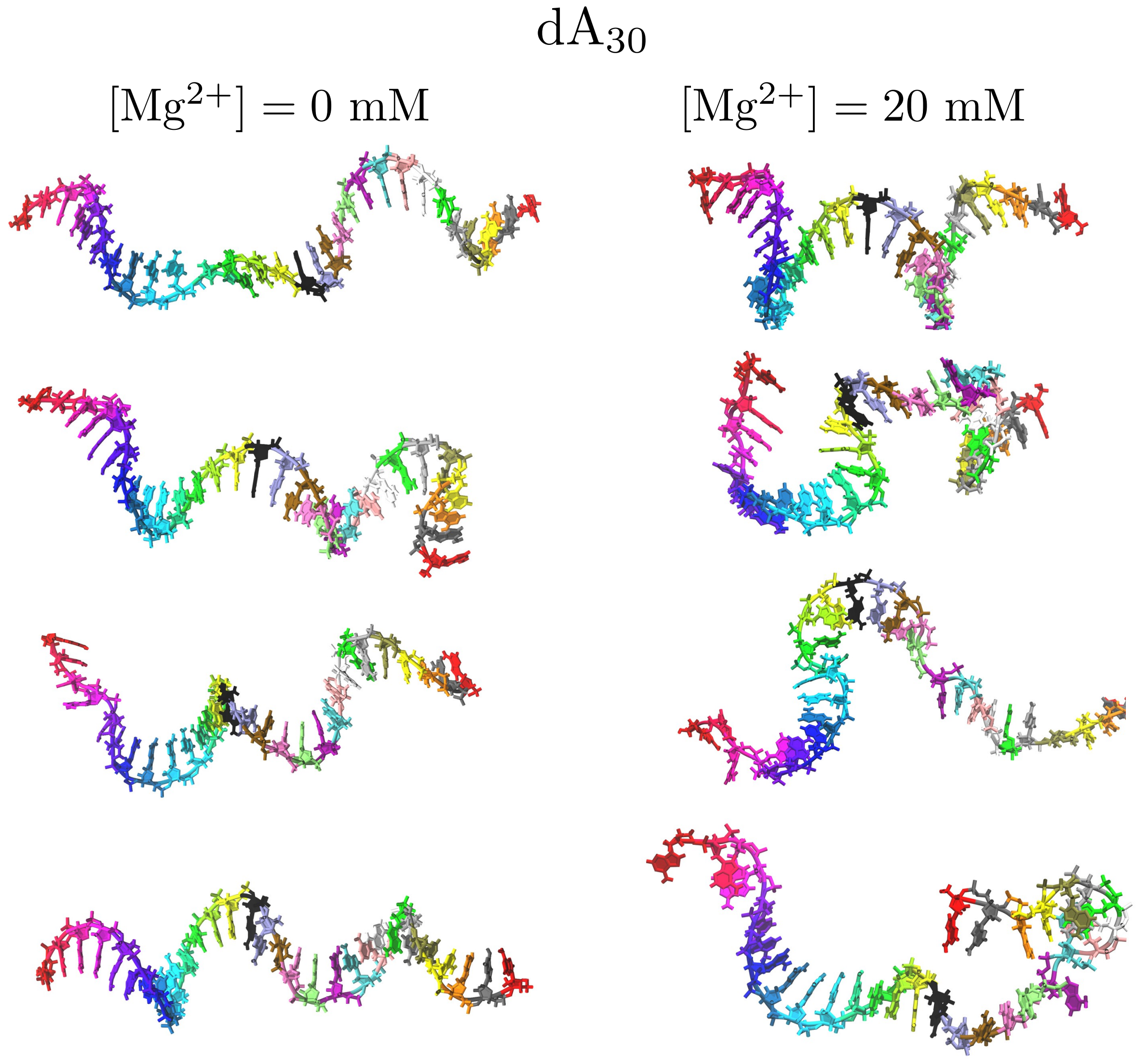


Figure S6: Same as Figure S5, except the results are shown for dA$_{30}$. The structures have considerable helical order.

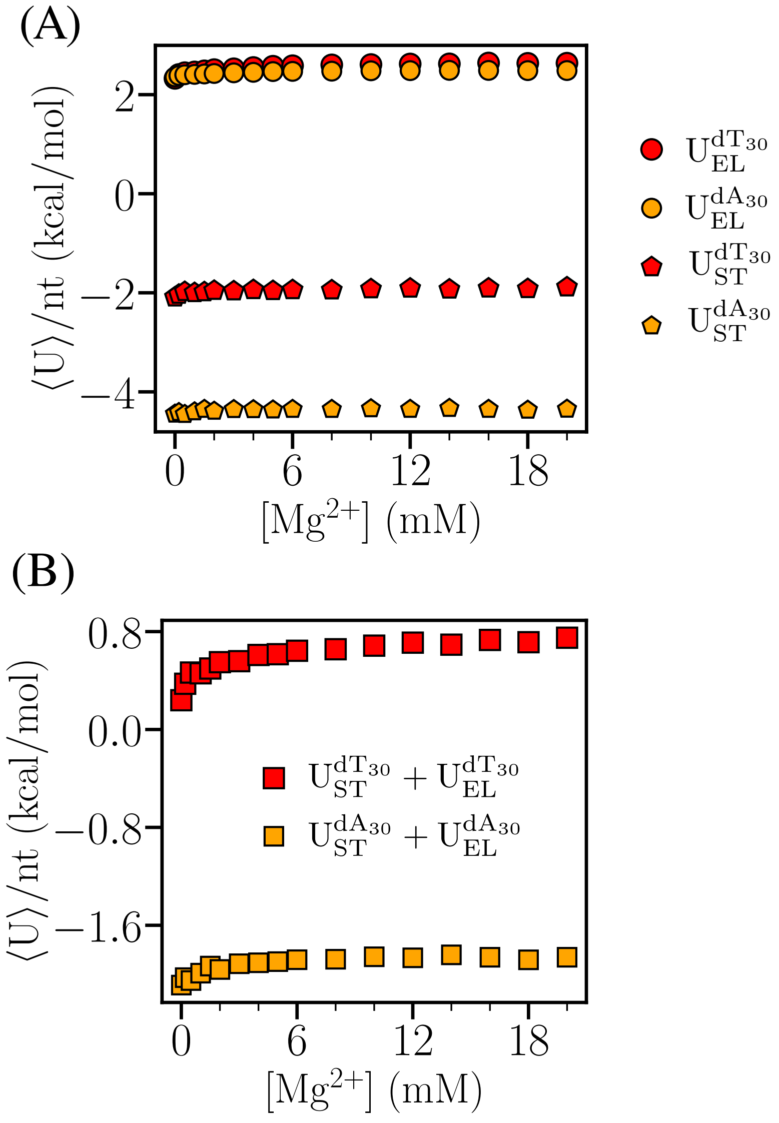


Figure S7: (A) Stacking ($U_{ST}$) and electrostatic ($U_{EL}$) components of the potential energy per nucleotide as a function of [$Mg^{2+}$] for $dT_{30}$ and $dA_{30}$. (B) Sum of $U_{ST}$ and $U_{EL}$ as a function of [$Mg^{2+}$].

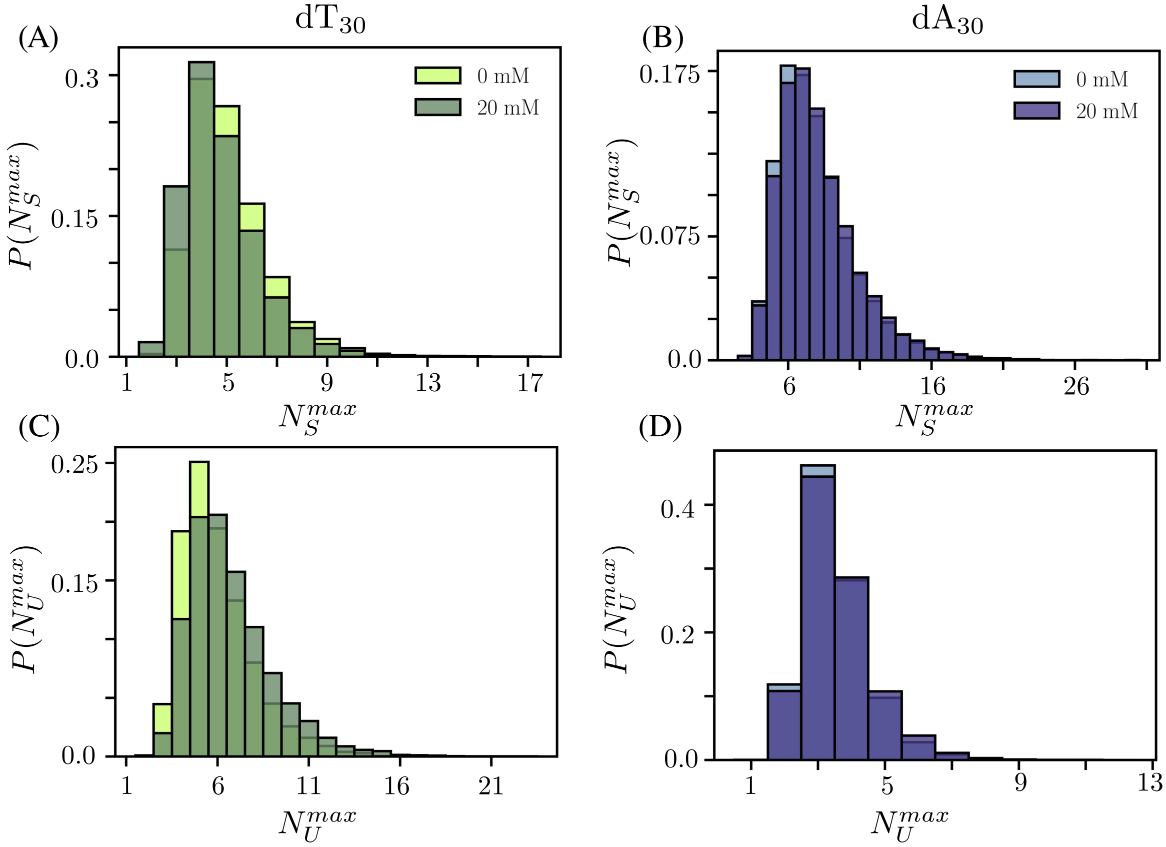


Figure S8: [$Mg^{2+}$] dependent changes in the distributions of the longest stretches of persistent stacks ($N_S^{max}$) and longest stretches of the unstacked regions ($N_U^{max}$) in dT$_{30}$ and dA$_{30}$. Differences are more pronounced in dT$_{30}$

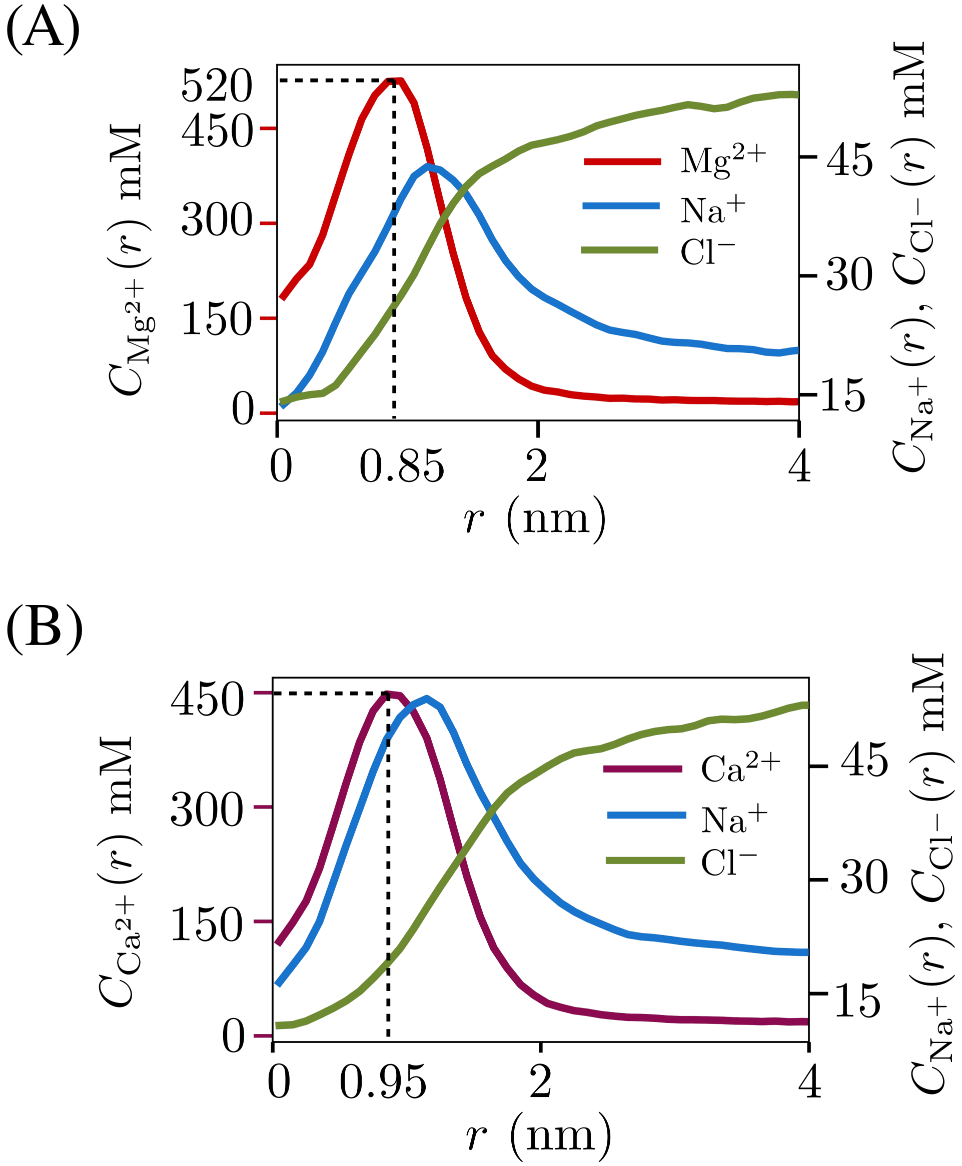


Figure S9: (A) Spatial dependence of ion concentrations, $C_{Mg^{2+}}(r)$, $C_{Na^+}(r)$ and $C_{Cl^-}(r)$, near dsDNA at $[Mg^{2+}] = 20$ mM. The scale on the right is for concentrations for $Na^+$ (blue) and $Cl^-$ (green). $r$ is the radial distance of an ion, measured from the principal axis of dsDNA. (B) Same as (A), except the results are for $[Ca^{2+}]$.

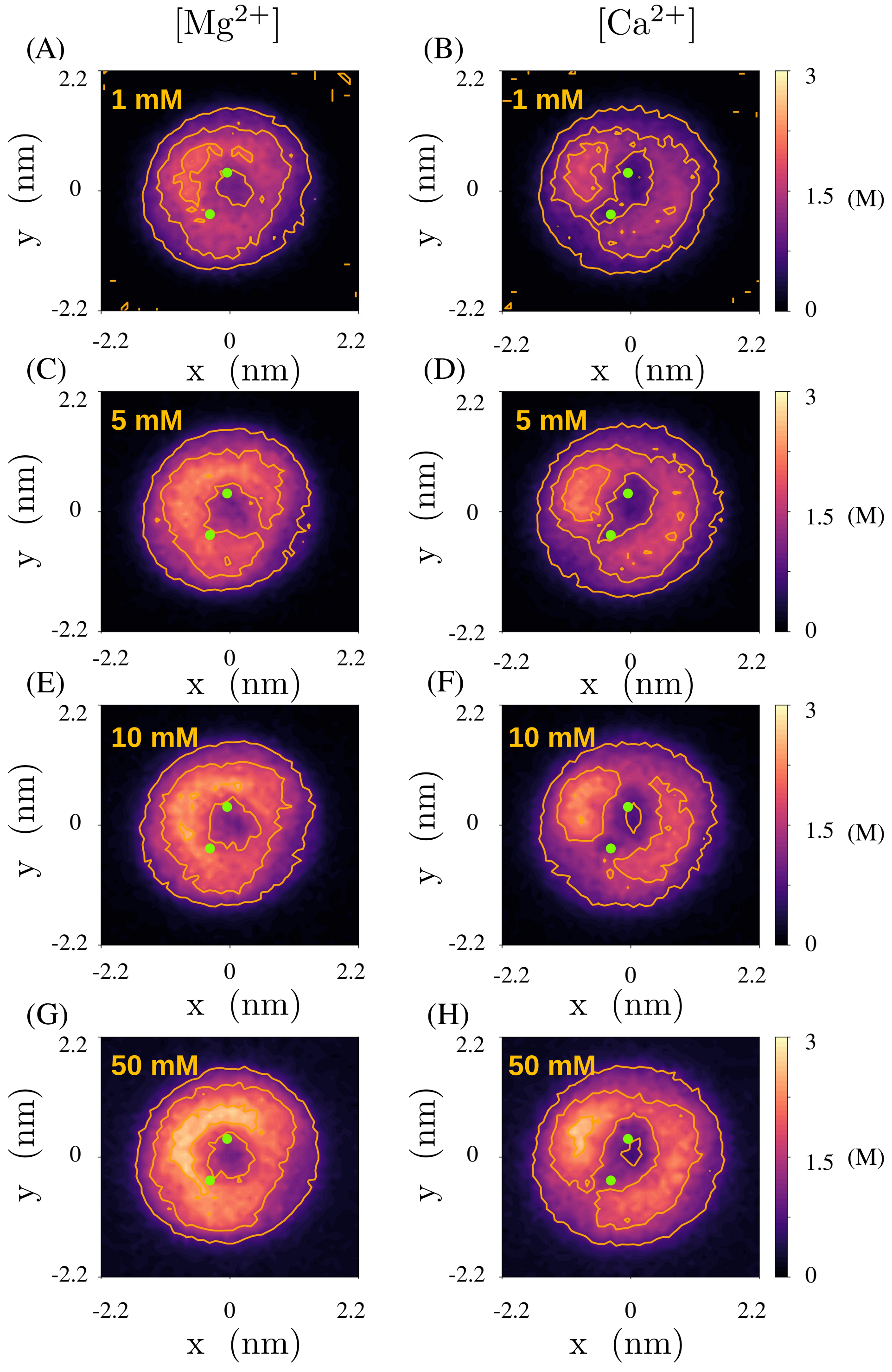


Figure S10: 2D untwisted densities of $Mg^{2+}$ and $Ca^{2+}$ ions projected onto an averaged base-pair plane of the 24 bp dsDNA. Averaged locations of the phosphate groups are highlighted in green.

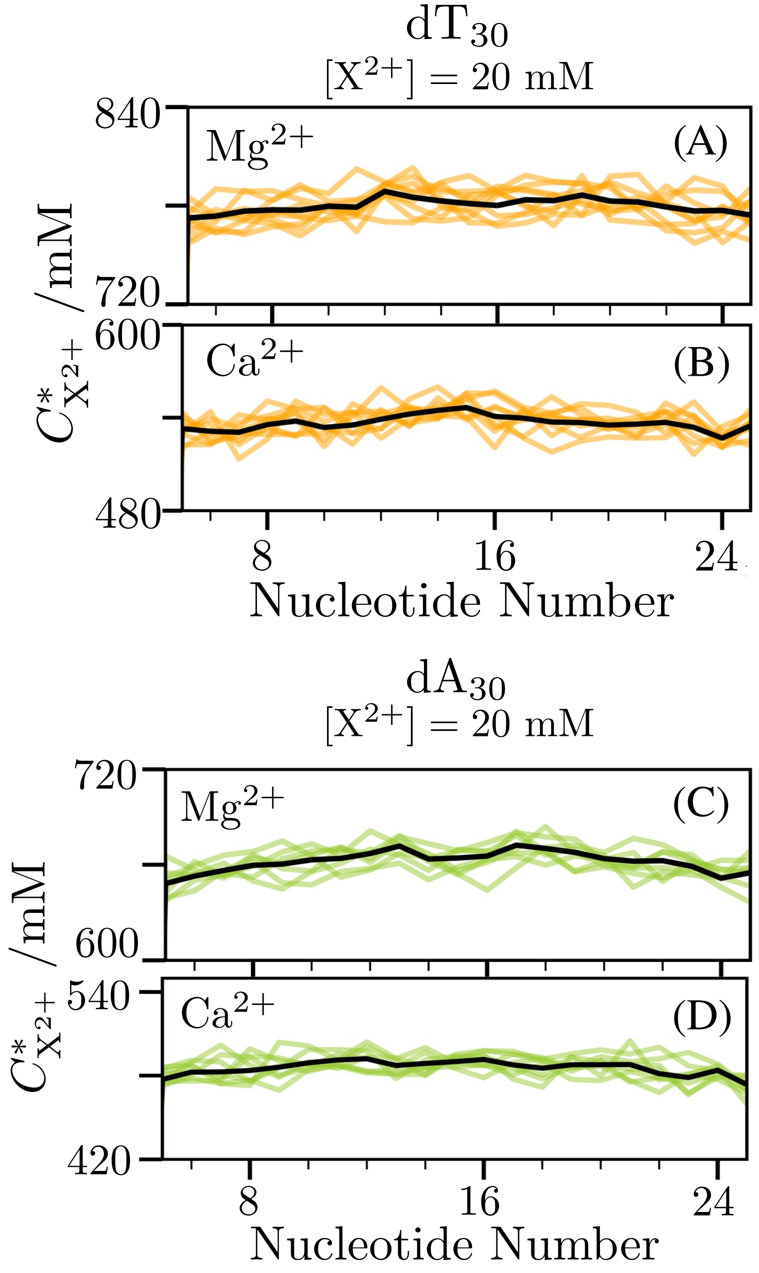


Figure S11: (A) Averaged local concentration of $Mg^{2+}$, $C^*_{Mg^{2+}}$, within a spherical shell centering the phosphate groups of $dT_{30}$ with inner radius $r_P$ and outer radius, $l_B$, the Bjerrum length. The black line is the globally averaged concentration profile using at least eight independent trajectories whereas lines in light color represent averaged concentrations corresponding to individual trajectories. (B) Same as (A), except the results are for $[Ca^{2+}]$. (C) Same as (A), except the results are for $dA_{30}$ sequence. (D) Same as (C), except the results are for $[Ca^{2+}]$.

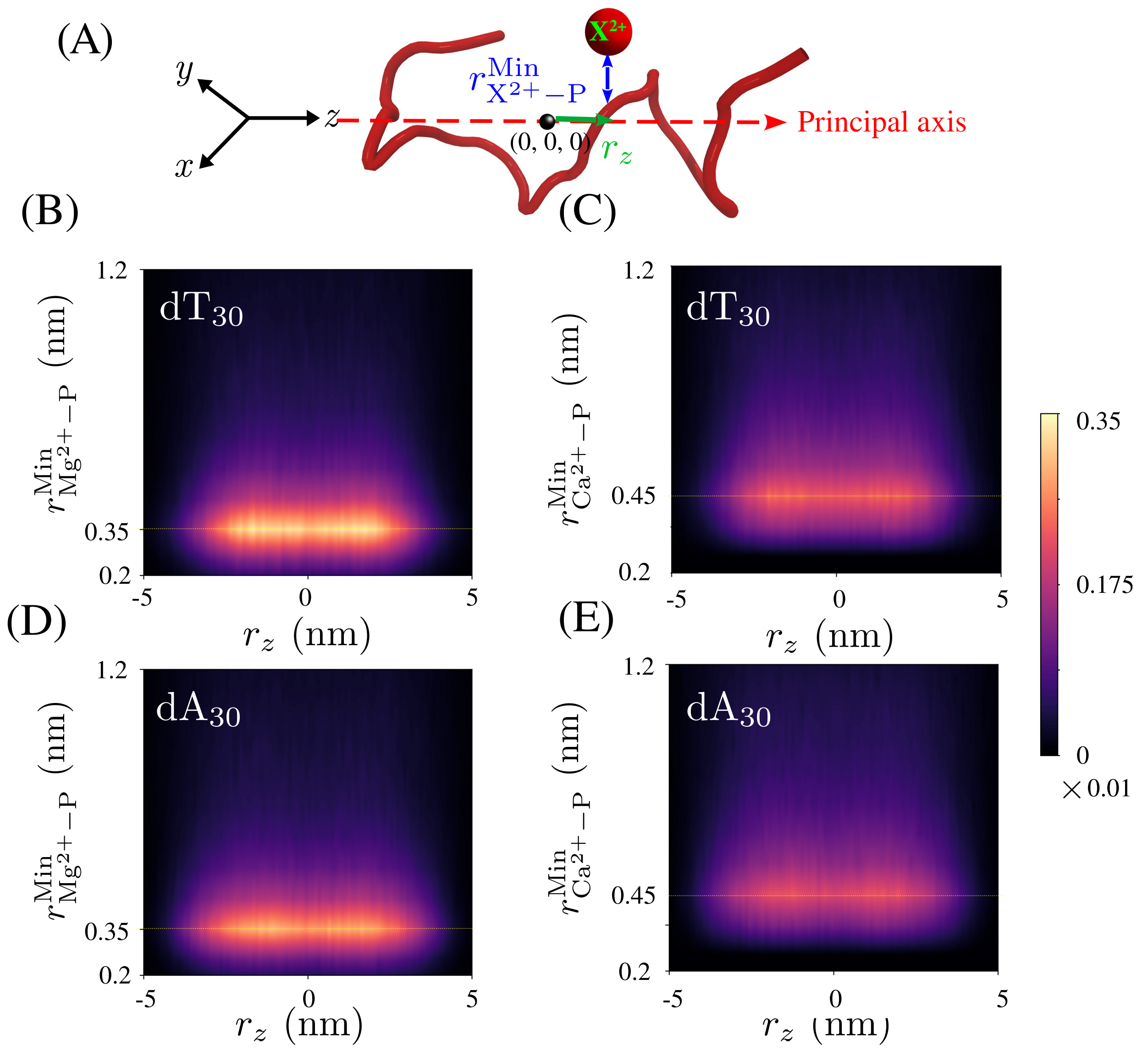


Figure S12: **Two-dimensional spatial distribution of divalent cations around ssDNA.** (A) The variables used in calculating the distribution are $r^{\mathrm{Min}}_{\mathrm{X^{2+}-P}}$ and $r_z$, using a typical ssDNA construct. Major principal axis (red dashed line) is aligned along the $z$ axis. Ion $X^{2+}$ is in red. $r^{\mathrm{Min}}_{\mathrm{X^{2+}-P}}$ is the distance of closest approach between phosphate and $X^{2+}$, and $r_z$ is the $z$-component of distance of $X^{2+}$ from the center of the major principal axis of ssDNA. (B) Joint distribution function of $r_z$ and $r^{\mathrm{Min}}_{\mathrm{X^{2+}-P}}$ for $dT_{30}$ sequence in $[Mg^{2+}] =$ 20 mM and in 20 mM NaCl solution. (C) Same as (B), except results are for $Ca^{2+}$ ions. (D) Same as (B), except results correspond to $dA_{30}$. (E) Same as (C), except results are for $dA_{30}$.

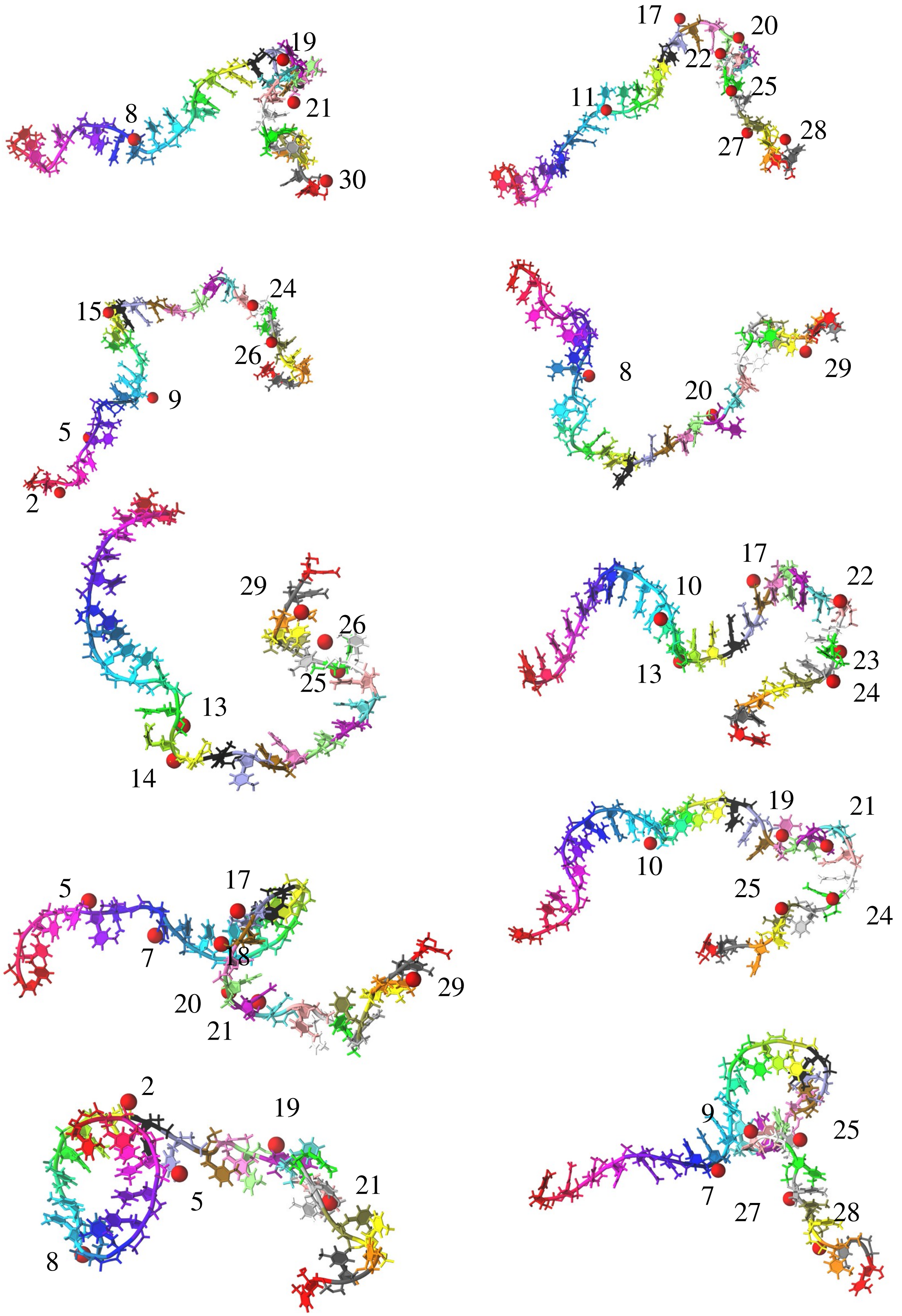


Figure S13: Direct interaction of $Mg^{2+}$ ions (in red) with the phosphate groups in $dT_{30}$. The concentration of $Mg^{2+}$ is 20 mM. There is considerable structural heterogeneity in the conformations.

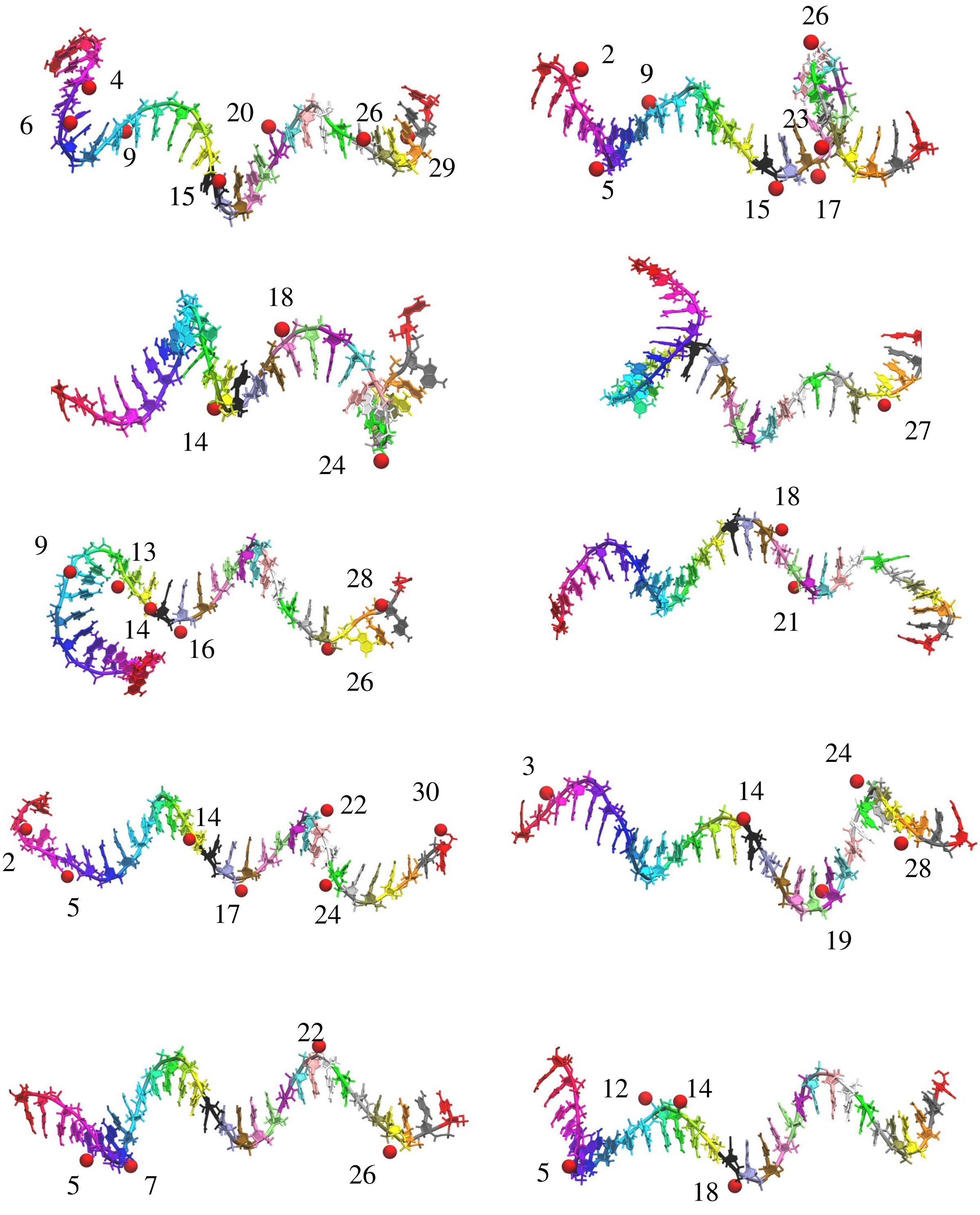


Figure S14: Same as Figure S13, except results are shown for $dA_{30}$. Preponderance of helical order is evident.

Table S1: Parameters for excluded volume and electrostatic interactions

| | $R_i$ nm | $\epsilon_i$ kcal/mol | $q_i$ e |
|---|---|---|---|
| P | 0.21 | 0.2 | -1 |
| S | 0.29 | 0.2 | 0 |
| A | 0.28 | 0.2 | 0 |
| G | 0.3 | 0.2 | 0 |
| T | 0.27 | 0.2 | 0 |
| C | 0.27 | 0.2 | 0 |
| $Mg^{2+}$ | 0.08 | 0.9 | 2 |
| $Ca^{2+}$ | 0.17 | 0.5 | 2 |
| $Na^{+}$ | 0.19 | 0.00277 | 1 |
| $Cl^{-}$ | 0.19 | 0.3 | -1 |

Table S2: Flory prefactor $R_0$ (Eq. 10 in the main text) in [$Mg^{2+}$]

| [$Mg^{2+}$] mM | $R_0$ ($dT_{30}$) nm | $R_0$($dA_{30}$) nm |
|---|---|---|
| 0 | 0.69 | 0.53 |
| 0.5 | 0.71 | 0.55 |
| 1.0 | 0.74 | 0.56 |
| 2.0 | 0.75 | 0.57 |
| 5.0 | 0.77 | 0.59 |
| 10.0 | 0.79 | 0.59 |
| 20.0 | 0.79 | 0.61 |

Table S3: Flory prefactor $R_0$ (Eq. 10 in the main text) in [$Ca^{2+}$]

| [$Ca^{2+}$] mM | $R_0$ ($dT_{30}$) nm | $R_0$($dA_{30}$) nm |
|---|---|---|
| 0 | 0.69 | 0.53 |
| 0.5 | 0.72 | 0.54 |
| 1.0 | 0.73 | 0.55 |
| 2.0 | 0.74 | 0.57 |
| 5.0 | 0.76 | 0.56 |
| 10.0 | 0.79 | 0.58 |
| 20.0 | 0.78 | 0.59 |

### References


(1) Hori, N. TIS2AA. 2017; `https://github.com/naotohori/TIS2AA/tree/v1.0`.

(2) Pérez, A.; Marchán, I.; Svozil, D.; Sponer, J.; Cheatham, T. E.; Laughton, C. A.; Orozco, M. Refinement of the AMBER force field for nucleic acids: Improving the description of $\alpha/\gamma$ conformers. *Biophys. J.* **2007**, *92*, 3817–3829.

(3) Zgarbová, M.; Sponer, J.; Otyepka, M.; Cheatham, T. E.; Galindo-Murillo, R.; Jurecka, P. Refinement of the sugar–phosphate backbone torsion beta for AMBER force fields improves the description of Z-and B-DNA. *J. Chem. Theory Comput.* **2015**, *11*, 5723–5736.

(4) Svergun, D.; Barberato, C.; Koch, M. H. CRYSOL - A program to evaluate X-ray solution scattering of biological macromolecules from atomic coordinates. *J. Appl. Crystallogr.* **1995**, *28*, 768–773.

(5) Schmitz, W. International Tables for X-ray Crystallography, Vol. IV, The Kynoch Press, Birmingham, England, 1974. *Kristall und Technik* **1975**, *10*, K120–K120.
